# DNA-SIP reveals salinity-associated niche differentiation of potentially active methanogens in mangrove soils

**DOI:** 10.64898/2026.07.05.736568

**Authors:** Ya-Wei Zeng, Yo-Jin Shiau

## Abstract

Mangrove forests are major blue carbon ecosystems but are often characterized by low surface methane (CH_4_) emissions. Such low emissions, however, do not necessarily indicate weak methanogenesis, because CH_4_ production may be offset by internal CH_4_ consumption before reaching the atmosphere. Although previous community, genomic, and transcriptomic studies have implicated methylotrophic methanogenesis in mangrove sediments, direct taxon-resolved evidence linking methylated carbon assimilation to potentially active methanogens remains limited. Here, we combined methanogenic activity assays, DNA stable isotope probing (DNA-SIP), *mcrA* and 16S rRNA gene analyses, and phylogenetic comparisons to identify potentially active methanogens across saline-influenced mangrove soils.

The results showed that CH_4_ production potentials were consistently dominated by methylotrophic pathways (1.86-2.78 μg CH_4_ g^-1^ soil hr^-1^) across all sites. DNA-SIP, together with consistent community patterns in fresh soils, indicated the potential activity of methylotrophic and mixotrophic methanogens under saline conditions. *Methanolobus*-affiliated methanogens were associated with salinity, Na^+^, Cl^−^, and NH ^+^, whereas *Methanosarcina* and unclassified *Methanosarcinaceae* were linked to soil soluble organic carbon availability and water content, indicating niche differentiation among active methanogenic groups. Phylogenetic analyses incorporating reference sequences from diverse environments further showed that potentially active mangrove methanogens were dominated by saline-associated lineages.

Together with our previous methanotrophic evidence from the same sites, these findings suggest that low CH_4_ emissions from mangrove blue carbon ecosystems can mask substantial internal CH_4_ cycling sustained by active methanogenesis and CH_4_ consumption.

## 1. Introduction

Climate change driven by increasing concentrations of greenhouse gases remains a major global concern. Among these gases, methane (CH_4_) plays a disproportionate role in radiative forcing relative to its atmospheric concentration, with a global warming potential substantially higher than that of carbon dioxide over a 100-year time horizon (IPCC, 2021). Wetlands are the dominant natural source of atmospheric CH_4_, and microbial CH_4_ production under anaerobic conditions strongly influences whether these ecosystems function as net carbon sinks or sources.

Mangrove forests have attracted increasing attention as important blue carbon ecosystems because of their high primary productivity and long-term carbon storage capacity (Alongi, 2014; Donato et al., 2011; Kauffman & Bhomia, 2017). In contrast to many freshwater wetlands, mangrove ecosystems are commonly characterized by relatively low surface CH_4_ emissions, a feature that has contributed to their perception as efficient natural carbon sinks.

A fundamental distinction between mangrove ecosystems and terrestrial freshwater wetlands lies in their geochemical background. Mangrove soils are regularly influenced by seawater, resulting in elevated salinity and high sulfate (SO_4_^2-^) concentrations compared to freshwater systems. These conditions have long been considered unfavorable for methanogenesis, as SO_4_^2-^ reduction is often assumed to outcompete CH_4_ production for shared substrates under saline conditions (Kuivila et al., 1989; Lovley & Klug, 1986; Schönheit et al., 1982; Shiau et al., 2016). As a result, mangrove soils are commonly regarded as systems where methanogenesis is strongly constrained relative to freshwater wetlands.

However, interpretations of low CH_4_ efflux in mangrove ecosystems are largely based on surface flux measurements and do not directly resolve subsurface CH_4_ cycling processes. In several coastal ecosystems, including the mangrove forests investigated in this study, previous work has consistently identified methane-oxidizing bacterial (MOB) communities dominated by methanotrophs affiliated with the family *Methylomonadaceae* (i.e. Type Ia methanotrophs) (Shiau et al., 2025; Shiau et al., 2020; Shiau et al., 2017a). These organisms are generally regarded as r-strategists adapted to environments with elevated CH_4_ availability (Ho et al., 2013; Knief, 2015; Shrestha et al., 2010). The widespread presence of such methanotrophs in systems exhibiting low net CH_4_ emissions suggests that CH_4_ production and CH_4_ emission may not be tightly coupled in mangrove soils, and that current interpretations based solely on surface fluxes may be incomplete.

Recent studies have increasingly recognized that methanogenesis in mangrove sediments is not necessarily suppressed under sulfate-rich conditions, but most evidence has been limited to omics-based observations and has not directly linked methanogenic potential to active functional groups. For example, community surveys and gene-based analyses have reported diverse methanogenic assemblages in mangrove soils and other coastal ecosystems, including frequent detection of methylotrophic and hydrogenotrophic methanogens (Cai et al., 2022; Euler et al., 2023; Yu et al., 2020; Zhang et al., 2020a; Zhang et al., 2025; Zhang et al., 2020b; Zhou et al., 2023). In addition, metatranscriptomic studies have further detected the expression of methanogenesis-related genes in mangrove sediments (Zhang et al., 2025; Zhang et al., 2020b), and substrate-amended incubations have provided important indications that methylated substrates can stimulate CH_4_ production in mangrove sediments (Dong et al., 2024). However, these approaches do not jointly link pathway-specific CH_4_ production rates with the methanogenic taxa responsible for substrate assimilation.

Together, these studies indicate that methanogenesis, particularly methylotrophic methanogenesis, may be more important in mangrove sediments than previously assumed.

However, community composition, gene abundance, transcript expression, and substrate-stimulation responses remain insufficient to directly identify which methanogenic taxa assimilate carbon into biomass. Direct evidence linking specific methanogenic taxa to substrate-derived carbon assimilation in mangrove soils therefore remains limited (Zhang et al., 2025).

DNA stable isotope probing (DNA-SIP) enables the identification of potentially active microorganisms through substrate incorporation and provides pathway-specific evidence of metabolic activity (Dunford & Neufeld, 2010; Neufeld et al., 2007). By linking labeled substrate incorporation to specific microbial taxa, DNA-SIP provides a direct complement to community, gene-based, and transcriptomic approaches. Previous applications of DNA-SIP have successfully identified potentially active methanogens in freshwater systems, such as rice paddies (Han et al., 2018; Lee et al., 2012; Xu et al., 2017), but comparable studies targeting methanogenic activity in mangrove forest soils remain scarce.

In this study, we applied DNA-SIP in combination with methanogenic activity assays, *mcrA* and 16S rRNA-based community analysis, and phylogenetic comparisons to investigate potentially active methanogens in mangrove forest soils along an estuarine salinity gradient, and to compare their phylogenetic relationships with methanogens reported from freshwater and saline environments across diverse geographic regions. The availability of the key substrate was also assessed to provide field context for the DNA-SIP incubations. The objectives were to determine which methanogenic pathways contribute to CH_4_ production under saline-influenced conditions, identify potentially active methanogens linked to substrate incorporation, and evaluate how salinity-related physicochemical factors shape methanogenic niche differentiation in mangrove soils. We hypothesized that low-emission, sulfate-rich mangrove soils retain substantial methanogenic potential, with non-acetoclastic pathways, particularly methylotrophic and hydrogenotrophic methanogenesis, sustaining CH_4_ production under saline conditions.

## 2. Materials and Methods

### 2.1 Site description and soil collection

Three mangrove forests with dominant plant species of *Kandelia obovata* were selected for this study in the Tamsui estuary (i.e. Tamsui river), Taipei, Taiwan (Figure 1). The first mangrove forest, Guandu, is located at the most upstream among the three study sites and is about 8 km away from the estuary (121° 27’ 51” E, 25° 06’ 55” N). The elevation of the mangrove forest is around 1.0 m above the sea level and the soil oxidation-reduction (i.e. redox) potential is around 48 mV.

**Figure 1.**
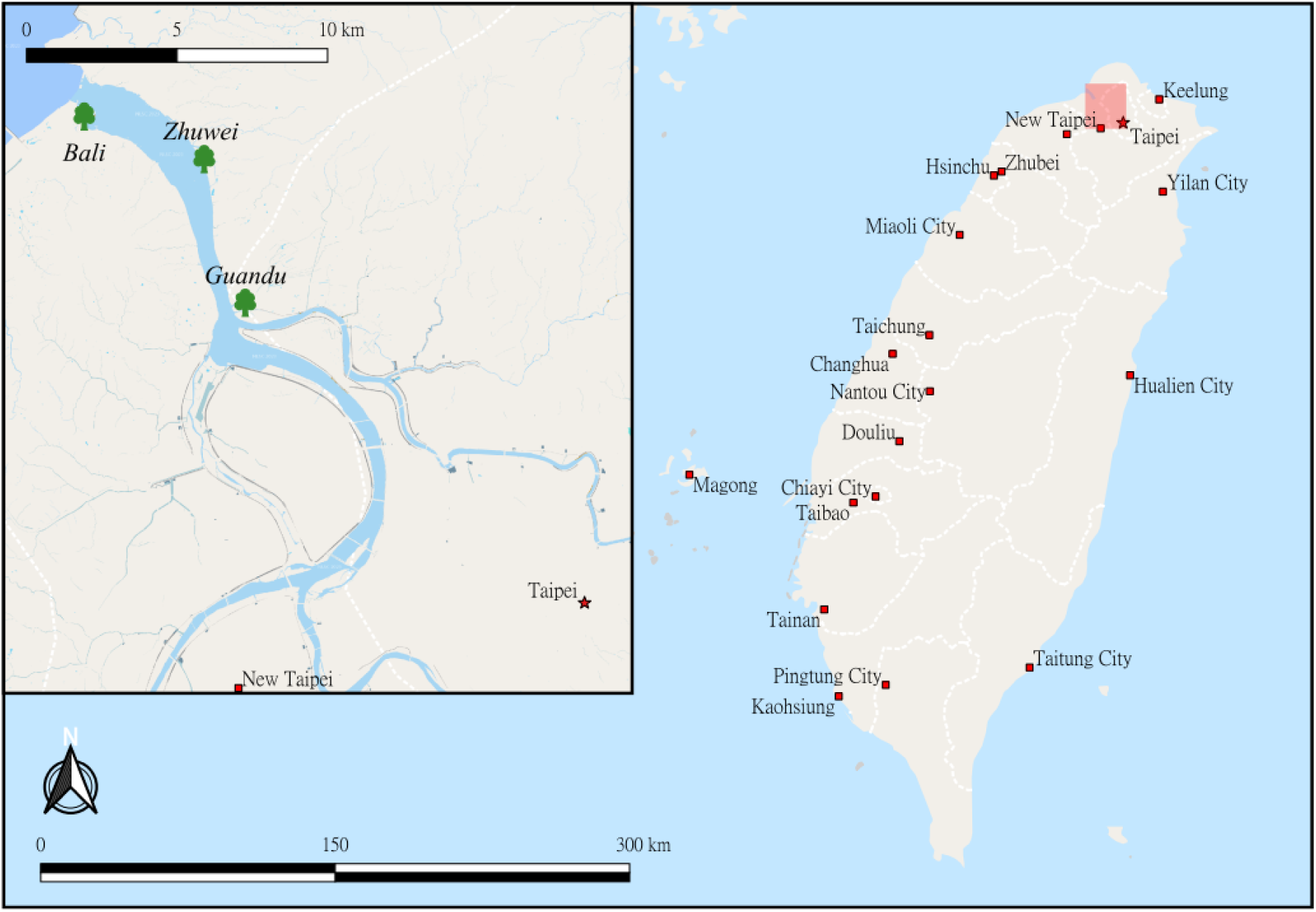
Locations of the mangrove forest study sites in the Tamsui River estuary, northern Taiwan. Sampling sites are indicated on the map.

The second mangrove forest, Zhuwei, is located about 5 km away from the estuary (121 ° 27’ 24” E, 25° 09’ 16” N). It has the highest elevation among the three sites, with an average elevation of 1.2 m above the sea level and a redox potential of 75 mV.

The third mangrove forest, Bali, is located at the most downstream and is at the mouth of the river facing to the sea (121° 25’ 00” E, 25° 09’ 56” N). The elevation of the site is around 0.8-0.9 m above the sea level and the overall soil redox potential is about -5 mV. The three study sites are all influenced by daily tidal cycles. Detailed site description can also be found in our previous studies (Shiau et al., 2020; Shiau et al., 2017a).

Three soil replicates were taken in each mangrove forest site on the same day during low tides in March 2022. To collect each replicate, 5 soil cores were taken with a steel soil sampler within the soil depth of 0-10 cm. Then, the 5 soil cores were mixed in a plastic zipper bag to form a composed replicate sample. The collected samples were immediately brought back to the laboratory and weighed for calculating the soil bulk density, and placed under 4 °C for further experiments.

In addition, soil pH and electrical conductivity (EC) were measured *in situ* with a portable multi-functional meter (WA-2017SD, Lutron, Taipei, Taiwan). Porewater salinity was measured with a refractometer (ATAGO Co., Tokyo, Japan).

### 2.2 Soil methanogenic activity determination

To analyze the methanogenic activity of mangrove soils, nine soil microcosms were prepared from each replicate sample in an anaerobic workstation (BACTRONEZ, Sheldon Manufacturing, Cornelius, USA). For each soil microcosm, about 2 g of composed soil was weighed out from each replicate, placed in a 30 ml serum vials, plugged with an anaerobic butyl rubber stopper, and capped with an aluminum cap.

Then, the nine soil microcosms from one replicate were divided into three groups. The three groups of soil microcosms were respectively amended with 0.5 ml of 400 mM methanol, 0.5 ml of 66.7 mM glucose, or 1.5 ml of CO_2_ and H_2_ based on previous study (Xu et al., 2017), so that every microcosm contained about the same amount of C (i.e. about 1000 μg C g^-1^ soil). Glucose was used as a fermentable substrate rather than acetate to avoid potential pH shifts associated with acetate amendment (Xu et al., 2017). The use of sodium acetate was also avoided to prevent changes in ionic conditions.

The microcosms were incubated in the anaerobic workstation under 25 °C for 2 weeks. On the Day 7 and 14, the CH_4_ concentrations in the headspace of the microcosms were analyzed with a gas chromatograph with a thermal conductivity detector (GC-TCD) (9790II, Fuli Instruments, Zhejiang, China) and a ShinCarbon ST column (Restek Corporation, Bellefonte, PA, USA).

The CH_4_ producing potentials of each microcosm were calculated by the linear regression of the CH_4_ concentrations with time.

### 2.3 DNA-SIP incubation and gradient fractionation

When the soil methanogenic activity experiments revealed the major C source for methanogenesis in the studied mangrove forests, another six soil microcosms were prepared from each replicate sample in the anaerobic workstation.

The six microcosms from each replicate were separated into two groups. One group of the microcosms were injected with the same quantity of the major C source containing pure (99 atom%) ^13^C, while the other group were injected with the same amount (99 atom% ^12^C) C source. The microcosms were incubated in the anaerobic workstation under 25 °C. On the Day 6, 10, 14, and 19, the headspace of the microcosms was analyzed for CH_4_ concentrations with the GC-TCD.

About 0.8 g of the incubated soil from each microcosm was extracted for the genomic DNA with DNA extraction kits (PowerSoil® DNA isolation kit, Qiagen, Hilden, Düsseldorf, Germany) based on the manufacturer’s instructions. The extracted DNA for each microcosm was partitioned into 15 density gradient fractions with an ultracentrifuge (CP80WX, Himac, Hitachi, Ltd., Tokyo, Japan) and a vertical rotor (P65VT2, Himac, Hitachi, Ltd., Tokyo, Japan) under 190,000 × g and 20 °C for 63 hrs. Detailed fractionation technique summarized in previous study (Lu & Jia, 2013). A digital refractometer (DR201-95, KRUSS, Hamburg, Germany) was used to measure the buoyant density for fractions 2-14.

After the gradient fractionation, the abundance of alpha subunit of *mcr* genes (i.e. *mcrA*) was quantified by real-time quantitative polymerase chain reaction (qPCR) (CFX connect, Bio-Rad Laboratories, Inc., California, USA) with a primer pair (*mcrA qF*/*mcrA qR*) (West et al., 2012) and a qPCR reagent kit (SYBR® Green Supermix, Bio-Rad Laboratories, Inc., California, USA).

In addition, DNA was also extracted from the fresh mangrove soils for determining the *mcrA* genes with qPCR.

### 2.4 Next generation sequencing (NGS) of the 16S rRNA and *mcrA* genes

The DNA heavy fractions of each ^13^C incubated sample with the highest *mcrA* genes from DNA-SIP gradient fractionation were sequenced for both the 16S rRNA and *mcrA* fragments using the Illumina MiSeq platform to identify the active methanogenic communities. The adjacent fractions were sequenced if their relative *mcrA* copy numbers exceeded 50% in comparison to the fractions with the highest *mcrA* copies from each incubated soil sample.

The DNA libraries of the *mcrA* genes were prepared with a polymerase chain reaction (PCR) (MiniAmp, Thermo Fisher Scientific, Waltham, MA, USA) of the *mcrA* gene with the primer pair *mlas-mod-F/mcrA-rev-R* (Angel et al., 2012; Steinberg & Regan, 2008) and a PCR reagent mix (Taq Master Mix RED reagent kit, AMPLIQON, Odense M, Denmark). The DNA libraries of the 16S rRNA genes were prepared with universal primer pair 515F/806R (Caporaso et al., 2012).

In addition, the genomic DNA from all *in situ* soil samples were also sequenced with *mcrA* and 16S rRNA genes for comparison.

### 2.5 Soil physiochemical properties analyses

Soil cations and anions, including SO_4_^2-^, nitrate (NO_3_^-^), nitrite (NO_2_ ^-^), ammonium (NH_4_^+^), sodium (Na^+^), potassium (K^+^) and chloride (Cl^-^), and soil soluble organic carbon (S_b_OC) were analyzed with a hot-water extraction method based on previous studies with small modifications (Shiau et al., 2017b). Briefly, 2 g of soil samples were mixed with 10 ml of deionized water in 15 ml centrifuge tubes and water bathed at 70 °C for 18 hours. Then, the tubes were shaken at 150 rpm for 5 mins and centrifuged at 10,000 × g for 10 mins to collect the suspension.

The suspension was filtered with 0.45 μm filters and analyzed for the cations and anions with an ion chromatography (IC) (925 ECO IC, Metrohm, Herisau, Switzerland). The filtered suspension was also analyzed for S_b_OC with a TOC analyzer (Aurora OI TOC 1030W, O.I. Corporation, TX, USA).

Subsamples of the collected soil samples were air dried, ground and analyzed with an elemental analyzer (PerkinElmer 2400, PerkinElmer, Shelton, CT, USA) for the soil total organic carbon (TOC) and total nitrogen (TN).

To assess the availability of methylated substrates, methanol concentrations were measured by static headspace method following Qu et al. (2018), but without acidifying the samples to prevent labile organic matter degradation and subsequent abiotic methanol production. Briefly, 5.0 g of surface soil was placed into 30 mL serum vials, sealed with butyl rubber stoppers and aluminum crimp caps, and immediately injected with 3 mL saturated NaCl solution. Samples were incubated at 80 °C for 30 minutes to allow headspace equilibration. Methanol in the headspace was quantified using a gas chromatograph equipped with a flame ionization detector (GC-FID) (GC9720, Fuli Instruments, Nanjing, China) equipped and an RT-Q-BOND column (Restek Corporation, Bellefonte, PA, USA). Methanol concentrations were determined by interpolation from calibration curves (0–2400 μM) constructed using known methanol standards in quartz sand. All samples were analyzed within 6 hrs to minimize methanol degradation.

In addition, about 10 g of subsamples were weighed and used to calculate soil gravimetric water content (WC) based on their fresh and oven-dried weights. All analytical data were converted to dry weight basis.

### 2.6 Data processes and statistical analysis

The sequence data was assembled and calculated for the operational taxonomic units (OTUs) using Mothur version 1.47 (Schloss et al., 2009). The 16s rRNA sequences were classified with RDP V18 and Silva 138 database, and the *mcrA* sequences were classified with the database published by Yang et al. (2014). The relative abundance of methanotrophs was averaged for samples containing multiple heavy fractions. The 50 most abundant *mcrA* OTUs from this study were analyzed together with 149 reference *mcrA* sequences retrieved from the NCBI database to place the identified methanogens into a broader phylogenetic context. Similarly, the 55 most abundant methanogenic 16S OTUs were analyzed with 169 reference sequences using nucleotide sequences. Both phylogenetic trees were reconstructed using the maximum-likelihood method implemented in MEGA X (Kumar et al., 2018), with branch support estimated from 1000 bootstrap replicates, and then visualized and annotated using Evolview 3.0 (Subramanian et al., 2019).

Soil physicochemical properties and variations in methanogenic activity and gene abundances (16S rRNA and *mcrA* copies) were evaluated with one-way ANOVA. Both analyses were followed by Tukey’s honestly significant difference (HSD) post-hoc test using JMP 11.0 (SAS Inc., Cary, NC, USA). Prior to analysis, gene copy numbers were log-transformed to satisfy normality assumptions, whereas soil physicochemical properties were tested for normality using the Shapiro–Wilk test, with Box–Cox transformation applied when necessary. A significant level of 0.05 was set for all the statistical analyses.

To evaluate the metabolic potential and niche differentiation within the dominant methanogenic community, differential abundance analysis was performed on the top 50 most abundant OTUs using the ALDEx2 package (v1.42.0) (Fernandes et al., 2014) as recommended by Nearing et al. (2022). Raw sequence counts were used as input to generate 128 Monte Carlo samples from a Dirichlet distribution, followed by Centered Log-Ratio (CLR) transformation. Statistical significance was determined based on expected FDR-corrected *p*-values using the Benjamini-Hochberg procedure.

To further distinguish robust biological patterns from experimental noise, effect sizes were evaluated to ensure that the mean difference between sites exceeded the maximum dispersion observed within replicates. Effect size thresholds have been suggested in previous studies (e.g., >0.6) (Fernandes et al., 2014), while a more conservative threshold (>1) was applied in this study to enhance robustness. High-fidelity rendering of taxonomic labels was implemented via the ggtext package (v0.1.2) in ggplot2 (v4.0.1). All statistical analyses and visualizations were performed in R (v4.5.2) (RStudio Team, 2020).

To investigate the relationships between soil physicochemical properties and representative methanogen communities in the fresh mangrove soils, redundancy analysis (RDA) and Spearman correlation analysis were performed using the vegan (v2.7.2) and Hmisc (v5.2-3) packages, respectively (Oksanen et al., 2018). Prior to RDA, relative abundance data were Hellinger-transformed and environmental variables were standardized. The significance of the RDA model and environmental variables was assessed using permutation tests.

All the *mcrA* and 16S rRNA gene sequences obtained from MiSeq sequencing have been deposited in the NCBI under the BioProject number: PRJNA1445139.

## 3. Results

### 3.1 Soil physicochemical properties, methanogenic activity, and DNA-SIP labeling

The results of one-way ANOVA of soil physiochemical properties showed pH, TOC, TN, NO_3_^-^, NO_2_^-^, SO_4_^2-^, and salt-extractable methanol concentrations were similar among the three study mangrove forests (Table 1). However, the EC, salinity, NH_4_^+^, Na^+^, K^+^ and Cl^-^concentrations were found significantly the highest at the Zhuwei site. Meanwhile, the S_b_OC concentrations were the highest in the Guandu mangrove forest soils.

**Table 1.**
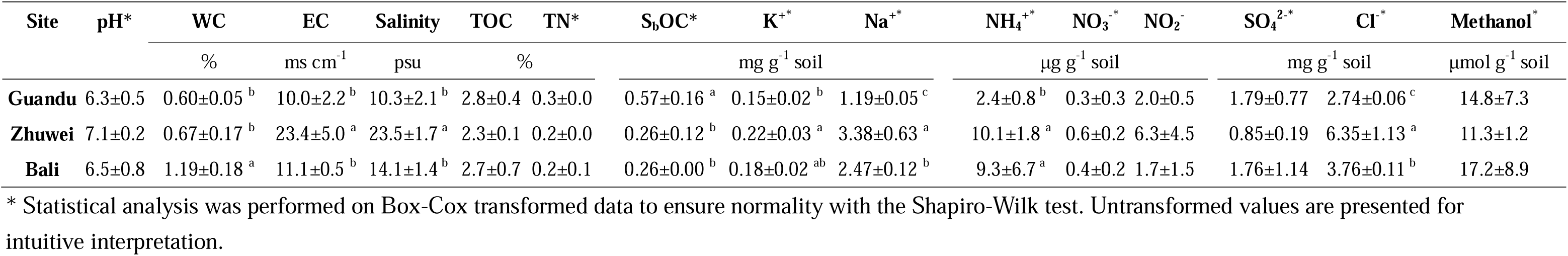
The physiochemical properties in the three studied mangrove forest soils at 0-10 cm depth.

The CH_4_ production potentials were 0.01 and 0.18 μg CH_4_ g^-1^ soil hr^-1^ under glucose and H_2_/CO_2_ amendments, respectively, and the potential was significantly higher (*p* < 0.05) under methanol amendment (2.32 μg CH_4_ g^-1^ soil hr^-1^) (Fig. 2a). The overall CH_4_ production potential was significantly higher in the Zhuwei and Guandu sites than the Bali site based on two-way ANOVA (*p* < 0.05).

**Figure 2.**
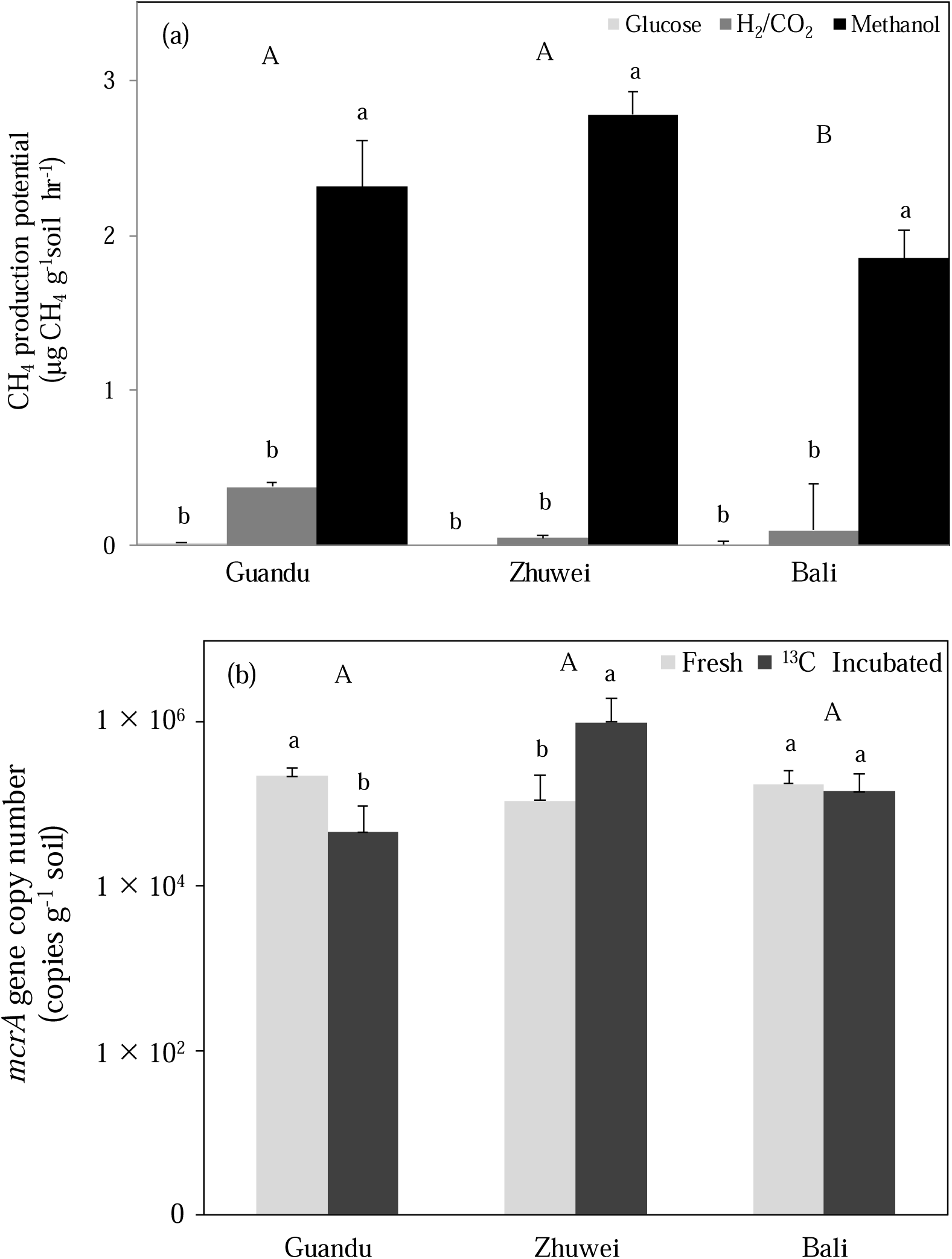
Methanogenic potential of mangrove soils under different substrate amendments and associated changes in methanogenic abundance. (a) Methane production potential in soils amended with glucose, H_2_/CO_2_, or methanol. (b) Abundance of methanogens quantified by *mcrA* gene copy numbers before and after ^13^C substrate incubation. Different uppercase letters indicate statistically significant differences among sites and lowercase letters indicate statistically significant differences among treatments based on two-way ANOVA and Tukey’s HSD test (*P* < 0.05).

Because the highest CH_4_ production potentials were observed with amended methanol in the studied mangrove forest soils, ^13^C methanol was used for DNA-SIP incubation. qPCR analysis of *mcrA* genes showed that methanogenic populations were similar among the fresh soils of the three mangrove forests (Fig. 2b). After DNA-SIP incubation, the *mcrA* copies significantly increased and were the highest in the Zhuwei mangrove soils. In contrast, *mcrA* copy numbers did not change statistically in the Bali mangrove soils, whereas they decreased and were lowest in the Guandu mangrove soils.

The qPCR analyses of the DNA-SIP fractions showed that the ^13^C heavy fractions were mainly concentrated in fractions 5 to 8 with buoyant densities ranges between 1.750-1.727 g ml^−1^. However, the ^12^C light fractions were mainly concentrated in fractions 10-11, with buoyant densities ranging from 1.710-1.702 g ml^−1^ (Fig. S1). The high relative abundance of *mcrA* copies in the heavy fractions of ^13^C incubated treatments indicated that the active methanogens were successfully labeled with ^13^C.

### 3.2 *mcrA*-based fresh and ^13^C-labeled methanogenic communities

High-throughput sequencing of *mcrA* genes yielded approximately 40,000–79,000 and 35,000–120,000 raw sequence reads from fresh soils and ^13^C-labeled heavy DNA fractions, respectively, across the three studied mangrove forests. The results revealed that *Methanolobus*, *Methanosarcina*, and unclassified methanogens affiliated with *Methanosarcinales* and *Methanosarcinaceae* were among the dominant methanogenic taxa in the fresh soils (Fig. 3a). In addition, hydrogenotrophic methanogens affiliated with the order *Methanomicrobiales* and the family *Methanomicrobiaceae* were consistently detected at low relative abundances in the fresh soils of all three mangrove forests.

**Figure 3.**
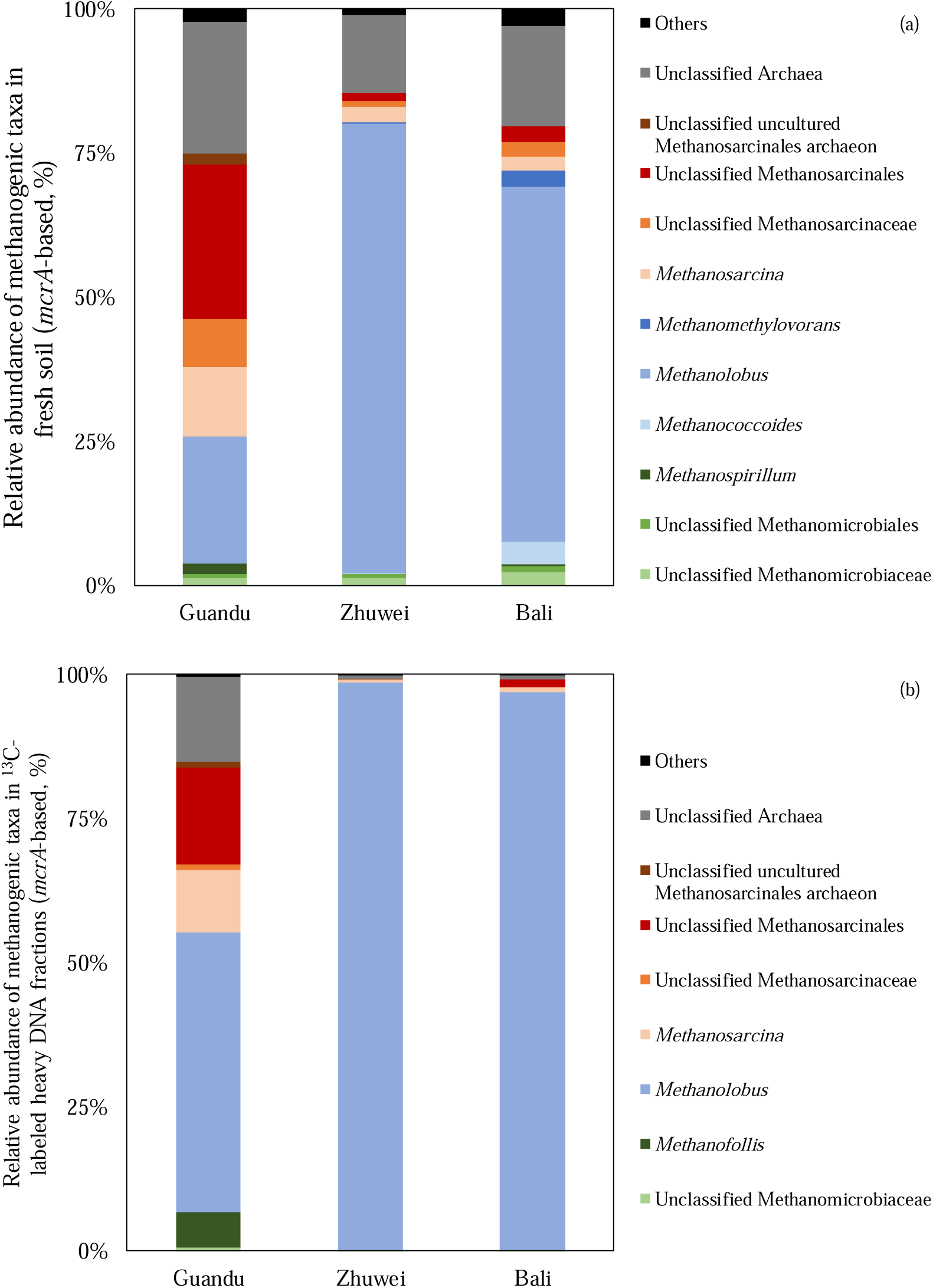
Community composition of methanogenic taxa based on *mcrA* gene amplicons in fresh soil (a) and ^13^C-labeled heavy DNA fractions (b) collected from mangrove forests in the Tamsui estuary, northern Taiwan. Colors represent methanogenic groups with different metabolic pathways (red, mixotrophic; blue, methylotrophic; green, hydrogenotrophic).

ALDEx2 analysis of the 50 most abundant *mcrA* OTUs showed that several methylotrophic methanogens, including OTU01, OTU05, OTU06, OTU07, OTU08, OTU29, OTU31 and OTU33 (unclassified *Methanolobus*), as well as OTU04 and OTU15 (*Methanolobus profundi*), made greater contributions in the Zhuwei and Bali mangrove soils than in the Guandu mangrove soils (FDR adjusted *p* < 0.05) (Fig. S2a; Table S1). In contrast, several mixotrophic methanogens did not differ significantly among the three sites (FDR adjusted *p* > 0.05), but their biological trends (i.e. absolute value of effect size > 1) suggested greater contributions in the Guandu mangrove soils, including OTU21, OTU37 and OTU48 (unclassified *Methanosarcinaceae*), and OTU09 (*Methanosarcina acetivorans*). Meanwhile, OTU32 (unclassified *Methanosarcinaceae*; mixotrophic), and OTU22 (unclassified McrA2 clone; unknown trophic type) showed greater contributions in the Bali mangrove soils. In addition, the hydrogenotrophic methanogens showed similar compositions among the three mangrove soils.

Results from the ^13^C-methanol labeled heavy DNA fractions showed that the potentially active methanogenic compositions were generally similar to those in the fresh soils across the three sites (Fig. 3b). Methylotrophic *Methanolobus* was dominant in the Zhuwei and Bali sites, whereas the Guandu soils were characterized by a more diverse assemblage of methanogens, including *Methanolobus*, *Methanofollis*, *Methanosarcina*, and unclassified methanogens affiliated with the order *Methanosarcinales*.

ALDEx2 analysis further identified OTU04 (*Methanolobus profundi*) and OTU07 (unclassified *Methanolobus*) as showing significantly greater relative contributions in the ^13^C-labeled Zhuwei soils than in the Guandu mangrove soils (FDR adjusted *p* < 0.05) (Fig. S2b; Table S2). Other potentially active methylotrophic methanogens, such as OTU05, OTU06, OTU13, OTU29, OTU31 and OTU33 (unclassified *Methanolobus*), as well as OTU12 (*Methanolobus profundi*), showed notable biological trends favoring the Zhuwei and Bali sites (FDR adjusted p > 0.05; absolute value of effect size > 1).

However, the ^13^C-labeled Guandu fractions were distinguished by the presence and greater relative contributions of potentially active methylotrophic OTU02 (*Methanolobus vulcani*), and several mixotrophic methanogens, including OTU09 (*Methanosarcina acetivorans*), OTU28 (unclassified *Methanosarcina*), OTU14 and OTU41 (unclassified *Methanosarcinaceae*), and OTU35 (unclassified *Methanosarcinales*) (FDR adjusted p > 0.05; absolute value of effect size > 1). In addition, potentially active OTU22 (unclassified McrA2 clone; unknown trophic type) showed greater relative contributions in the Bali mangrove soils, whereas OTU11 (unclassified *Archaea*; unknown trophic type) showed greater relative contributions in the Guandu mangrove soils.

### 3.3 16S rRNA gene-based fresh and ^13^C-labeled methanogenic communities

High-throughput sequencing of 16S rRNA genes yielded approximately 40–500 and 6,200–251,000 raw sequence reads affiliated with methanogenic taxa from fresh soils and ^13^C-labeled heavy DNA fractions, respectively, across the three mangrove forests.

Hydrogenotrophic *Methanobacterium*, methylotrophic *Methanolobus* and mixotrophic *Methanosarcina* were consistently detected as dominant groups in fresh soil samples from all three mangrove sites (Fig. 4a). However, results from the ^13^C-labeled heavy DNA fractions indicated that methylotrophic *Methanolobus* was the predominant potentially active methanogen across the three mangrove soils (Fig. 4b). In addition, mixotrophic *Methanosarcina* was also identified as potentially active in the Guandu mangrove soils.

**Figure 4.**
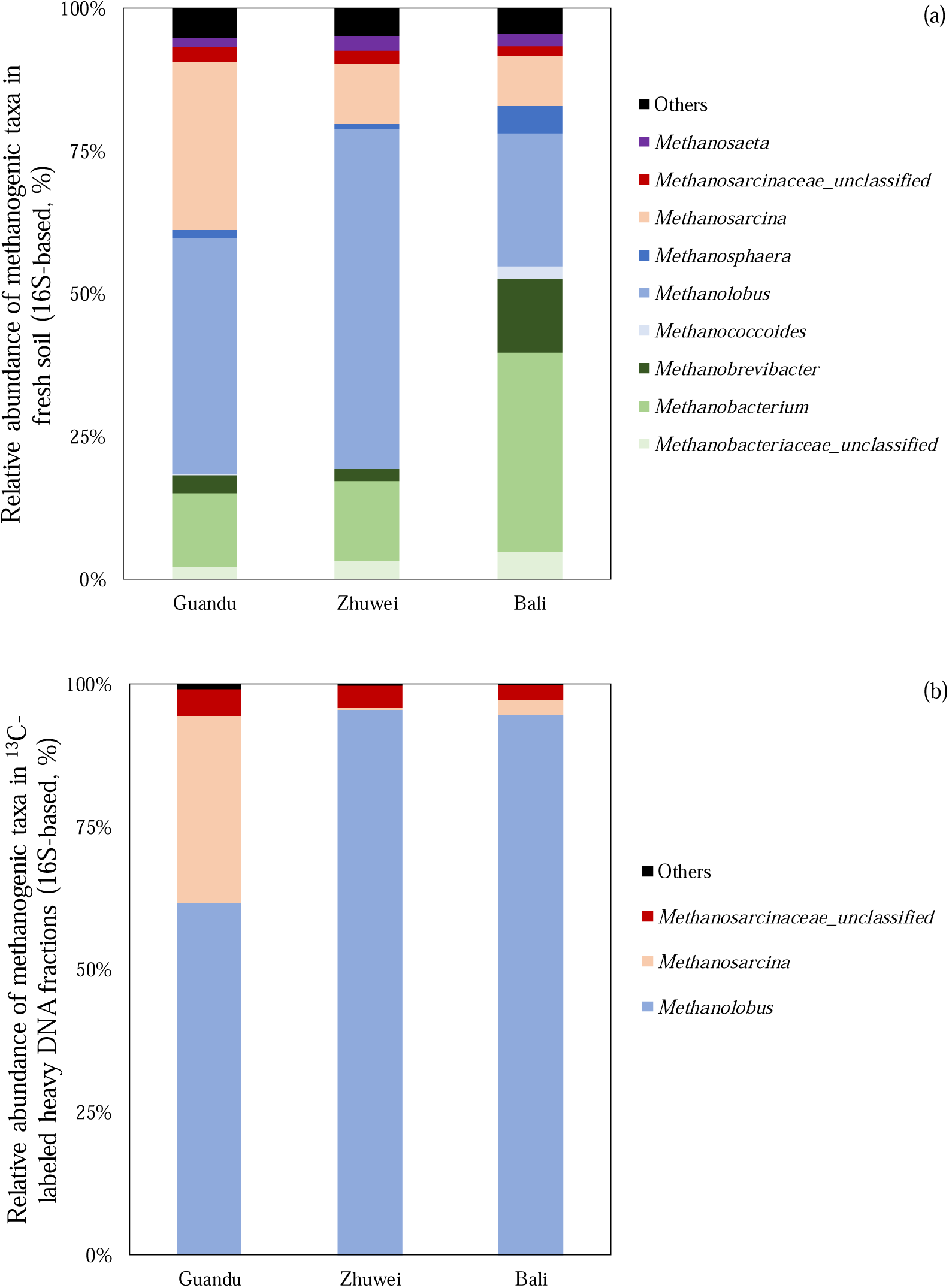
Community composition of methanogenic taxa based on 16S rRNA gene amplicons in fresh soil (a) and ^13^C-labeled heavy DNA fractions (b) collected from mangrove forests in the Tamsui estuary, northern Taiwan. Colors represent methanogenic groups with different metabolic pathways (red, mixotrophic; blue, methylotrophic; green, hydrogenotrophic; purple, acetoclastic).

### 3.4 Environmental and phylogenetic niches of potentially active methanogens

RDA integrating soil physicochemical properties showed that methylotrophic *Methanolobus* was associated with soil salinity and NH_4_^+^ concentrations, while other methylotrophic methanogens, such as *Methanococcoides* and *Methanomethylovorans*, were associated with soil SO_4_ ^2-^ concentrations (Fig. 5). In contrast, *Methanosarcina* and other unclassified mixotrophic methanogens belonging to the family *Methanosarcinaceae* were associated with soil S_b_OC concentration and soil WC. At the OTU level, RDA results showed consistent patterns, with methylotrophic methanogens related to seawater-associated factors (i.e. K^+^, Na^+^, Cl^-^, EC, and salinity), while mixotrophic methanogens were related to S_b_OC and WC (Fig. S3).

**Figure 5.**
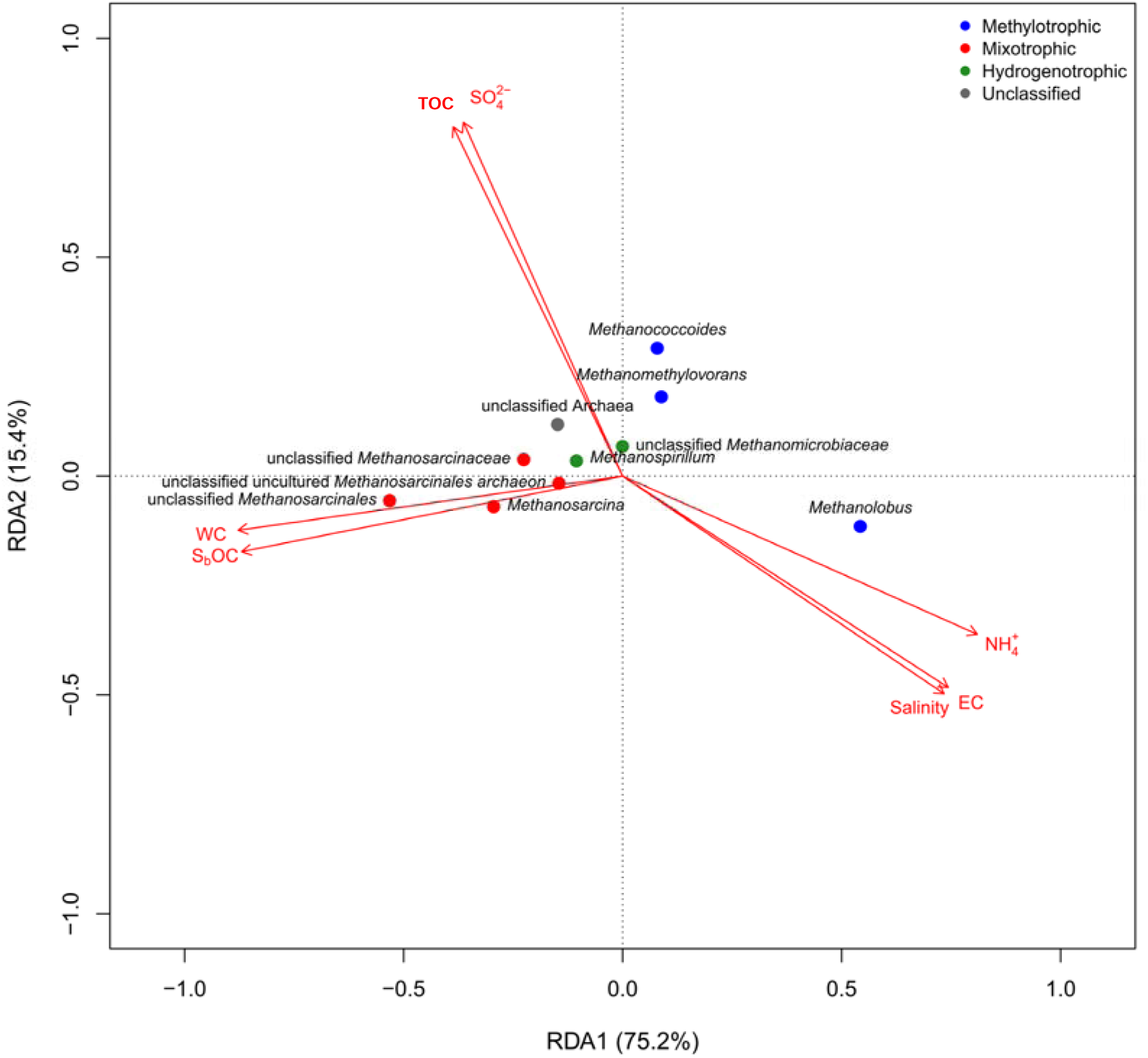
Redundancy analysis (RDA) showing the relationships between methanogenic taxa and soil physicochemical properties in field soils from the three mangrove forests. Methylotrophic methanogens are shown in blue, mixotrophic methanogens in red, hydrogenotrophic methanogens in green, and unclassified methanogens in grey. [S_b_OC: soluble organic carbon, NH_4_^+^: ammonium, SO_4_^2-^: sulfate, EC: electrical conductivity, WC: water content, TOC: total organic carbon]

Spearman correlation analysis further revealed relationships between methanogenic taxa of different trophic types and soil physicochemical properties at both the genus and OTU levels (Fig. 6; Fig. S4). *Methanolobus* was positively correlated to soil salinity, EC, NH_4_^+^, Na^+^ and Cl^-^ concentrations, whereas *Methanosarcina* was positively correlated to S_b_OC concentration and soil WC. In contrast, unclassified *Methanosarcinales* and unclassified *Methanosarcinaceae* were negatively correlated with salinity, EC, Na^+^, K^+^ and Cl^-^concentrations.

**Figure 6.**
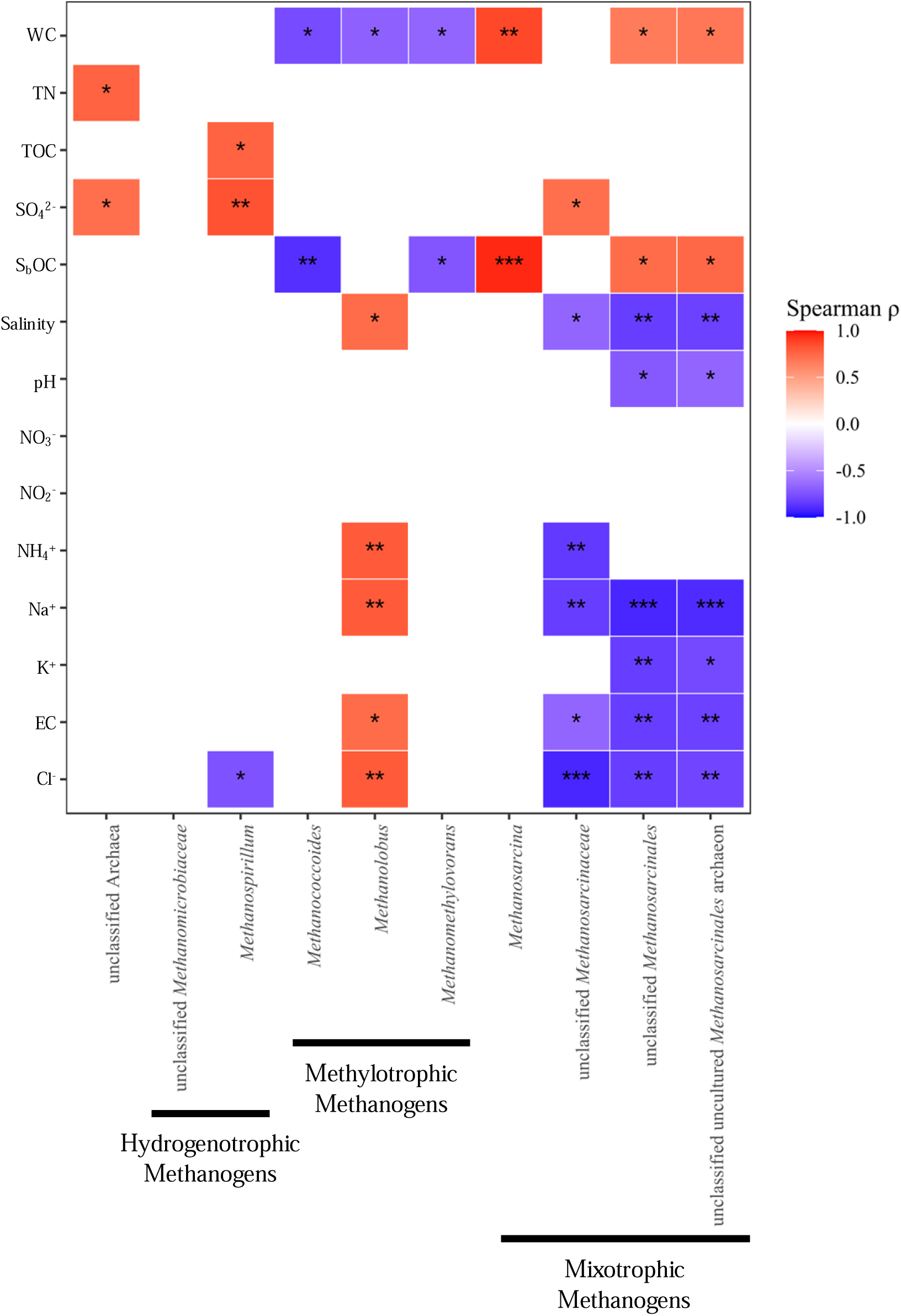
Spearman correlation analysis between soil physicochemical properties and methanogenic genera based on *mcrA* gene sequences in field soils from three mangrove forests along the estuarine gradient in northern Taiwan. (*: p<0.05; **: p<0.01; ***: p<0.001)

Phylogenetic analyses integrating representative *mcrA* OTUs from this study with 149 reference sequences, and methanogenic 16S rRNA OTUs with 169 reference sequences from the NCBI database, showed that the detected methanogens clustered within clades widely reported from coastal wetlands worldwide (Fig. 7; Figs. S5 and S6).

**Figure 7.**
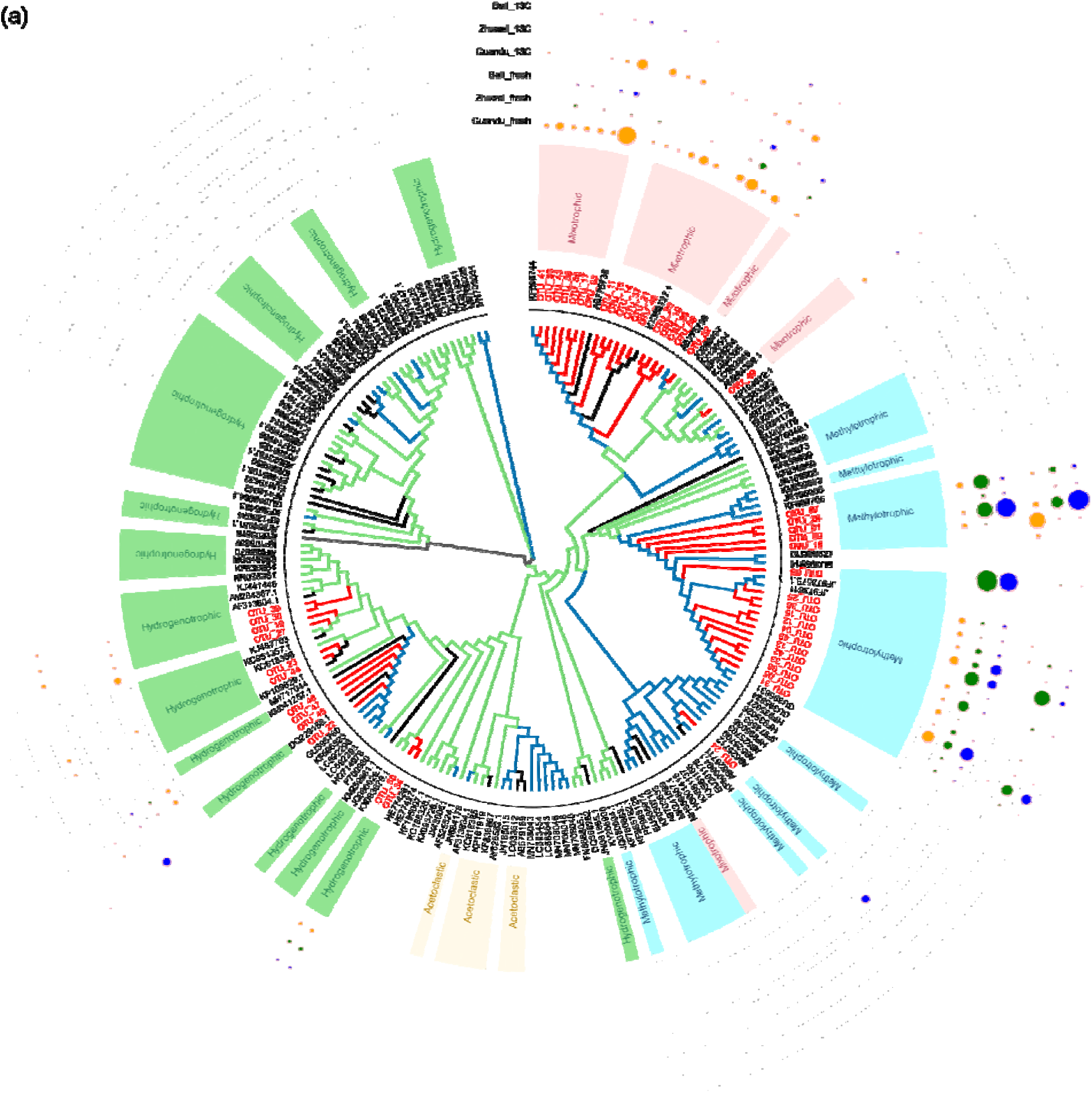
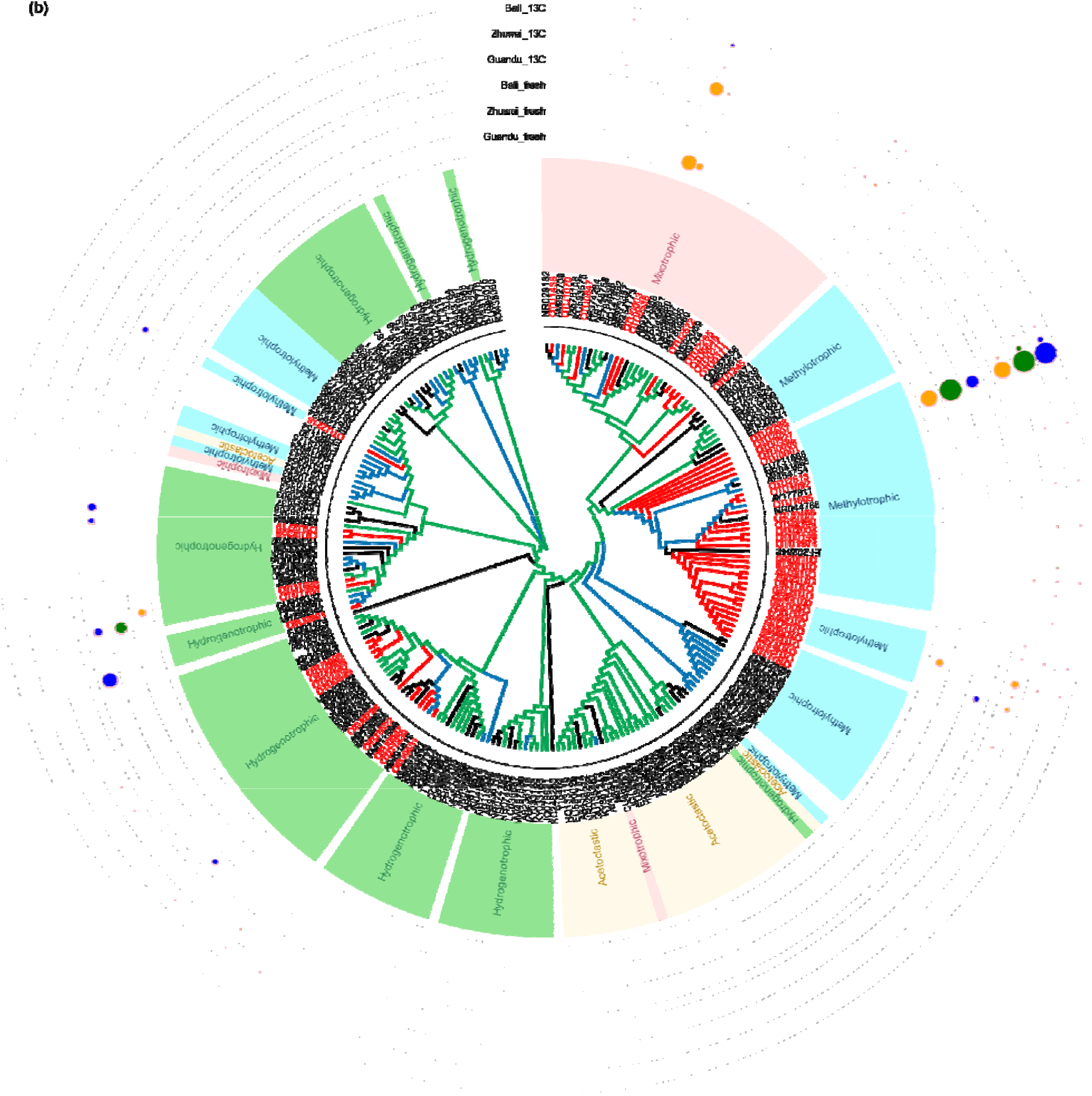
Phylogenetic placement of methanogenic lineages detected with *mcrA* (a) and 16S rRNA (b) genes in the studied mangrove soils. Maximum-likelihood phylogenetic trees were constructed using *mcrA* sequences including the 50 most abundant OTUs identified in this study together with 149 reference sequences retrieved from the NCBI database, and methanogenic 16S rRNA gene sequences including the 55 most abundant OTUs and 169 representative reference sequences. The reference sequences represent methanogenic lineages reported from diverse environments worldwide. Branch colors indicate the ecological origin of reference sequences, with blue branches representing saline-associated environments and green branches representing freshwater environments. OTUs detected in this study are highlighted with red labels and red branches. The circular dot plot indicates the relative abundance of OTUs across samples, with dot size proportional to the observed relative abundance.

## 4. Discussion

### 4.1 Salinity restructures potentially active methanogenic pathways beyond compositional patterns

The results of this study demonstrate that low-emission mangrove soils can retain substantial methanogenic activity, with CH_4_ production primarily supported by methylotrophic pathways. The integration of pathway-specific methanogenic activity assays with DNA-SIP linked CH_4_ production to substrate-assimilating methanogenic taxa, providing evidence beyond community composition, gene abundance, or transcript expression alone.

Salinity gradients are a defining feature of estuarine mangrove ecosystems and strongly influence anaerobic microbial metabolism by controlling ion availability, mediating substrate competition, and shaping redox energetics (Alongi, 1994; Kristensen et al., 2008; Tsola et al., 2021). Previous studies in coastal wetlands have consistently reported pronounced shifts in methanogenic community composition along salinity gradients, largely attributed to SO_4_ ^2-^availability, and typically characterized by a decline in acetoclastic methanogens alongside an increased prevalence of methylotrophic lineages under saline conditions (Tsola et al., 2021; Webster et al., 2015).

Consistent with these observations, our *mcrA*-based metabarcoding results revealed a clear reorganization of methanogenic assemblages across the estuarine system. This compositional shift occurred in parallel with distinct physicochemical differences across the three investigated mangrove forests. While bulk soil properties such as pH, TOC, TN, and SO_4_^2-^ concentrations were broadly comparable among sites, salinity-associated ions (Na^+^, K^+^, and Cl^-^) exhibited pronounced spatial variability rather than a simple downstream increase. Notably, the highest ion concentrations were observed at the midstream Zhuwei site, which may reflect local geomorphological and hydrological conditions. This site is characterized by a relatively elevated topographic position compared to the other mangrove forests, which may promote evaporative concentration and lead to localized salt accumulation.

Thus, these results suggest that restructuring of methanogenic communities in coastal ecosystems may not be governed by SO_4_^2-^ availability alone, but instead reflects the combined influence of salinity-associated ion composition and the spatial dynamics of labile organic carbon. Rather than following a simple monotonic gradient, these factors exhibited site-specific variability across the studied mangrove forests. RDA and Spearman correlation analyses further resolved these relationships, indicating positive associations between methylotrophic methanogens and salinity-driven physicochemical variables, whereas mixotrophic methanogens were more closely linked to S_b_OC availability and soil WC. Such patterns are consistent with previous studies showing that labile organic carbon inputs and hydrological conditions can influence methanogenic activity in coastal sediments (Chuang et al., 2016; Marinho et al., 2012; Torres-Alvarado et al., 2013).

Building upon the compositional patterns discussed above, methanogenic activity assays and DNA-SIP results further revealed substantial overlap between methanogenic assemblages detected in fresh soils and taxa enriched within the ^13^C-labeled DNA fractions. This suggests that many of the dominant methanogens observed *in situ* represent populations with high potential activity. In particular, methylotrophic *Methanolobus* dominated the labeled heavy DNA fractions in midstream and downstream mangrove soils, whereas upstream environments supported a broader contribution from mixotrophic *Methanosarcina*.

Meanwhile, the upstream Guandu site exhibited the lowest salinity-related ion concentrations and the highest S_b_OC levels, a readily metabolizable soil carbon fraction (Nishiyama et al., 2001), consistent with trends reported along the Tamsui River estuary (Wen et al., 2008). These environmental conditions may contribute to the higher diversity of methanogenic taxa observed at Guandu, including both methylotrophic and mixotrophic methanogens (Fig. S7). At the downstream Bali site, the highest salt-extractable methanol concentrations and a high relative abundance of methylotrophic methanogens in the ^13^C-labeled heavy fractions were observed, yet CH_4_ production potential was lowest among sites. Because *mcrA* gene abundance was broadly comparable across sites, this mismatch cannot be explained simply by methanogenic abundance or substrate availability. Instead, the significantly lower Shannon diversity of the active methanogenic community at Bali suggests that reduced community diversity, and potentially lower functional breadth, may be one factor constraining realized methanogenic activity even under apparently favorable substrate conditions.

At a finer resolution, OTU-level RDA and Spearman analyses further indicated that methanogens within the same trophic category generally shared similar environmental associations, while exhibiting minor OTU-specific variations (Fig. S5; S6). These variations suggest micro-scale niche partitioning within methylotrophic and mixotrophic guilds, without altering the overarching salinity-driven patterns across trophic categories.

Differences between methanogenic community profiles inferred from 16S rRNA and *mcrA* genes are likely attributable to marker specificity and sequencing depth, particularly given the low relative abundance of microbial functional groups in environmental samples, which can introduce variability in 16S-based profiles (Diwan et al., 2018). However, the enrichment of methanogenic populations in the ^13^C-labeled heavy DNA fractions substantially reduced this discrepancy, resulting in highly consistent patterns between 16S and *mcrA* datasets. This consistency further highlights the importance of marker choice and effective enrichment when assessing low-abundance functional guilds in complex soil communities.

### 4.2 Comparable CH_4_ production potentials between mangrove and freshwater wetland soils

For decades, mangrove ecosystems have been widely regarded as systems with high carbon sequestration capacity but low CH_4_ efflux, based on extensive field measurements and regional to global CH_4_ budget assessments (Alongi, 2014; Donato et al., 2011). This has commonly been attributed to the high SO_4_^2-^ concentrations in coastal environments, which favor sulfate-reducing bacteria (SRB) over methanogens (Lovley & Klug, 1983; Muyzer & Stams, 2008; Segarra et al., 2013).

However, previous studies have reported considerable CH_4_ production potentials across a range of coastal and brackish ecosystems, with values comparable to those observed in freshwater systems such as rice paddies, lakes, and freshwater wetlands (Table 2). When considered together with the results of the present study, these findings suggest that CH_4_ production potential in mangrove soils may not be inherently lower than that of freshwater wetland systems.

**Table 2.**
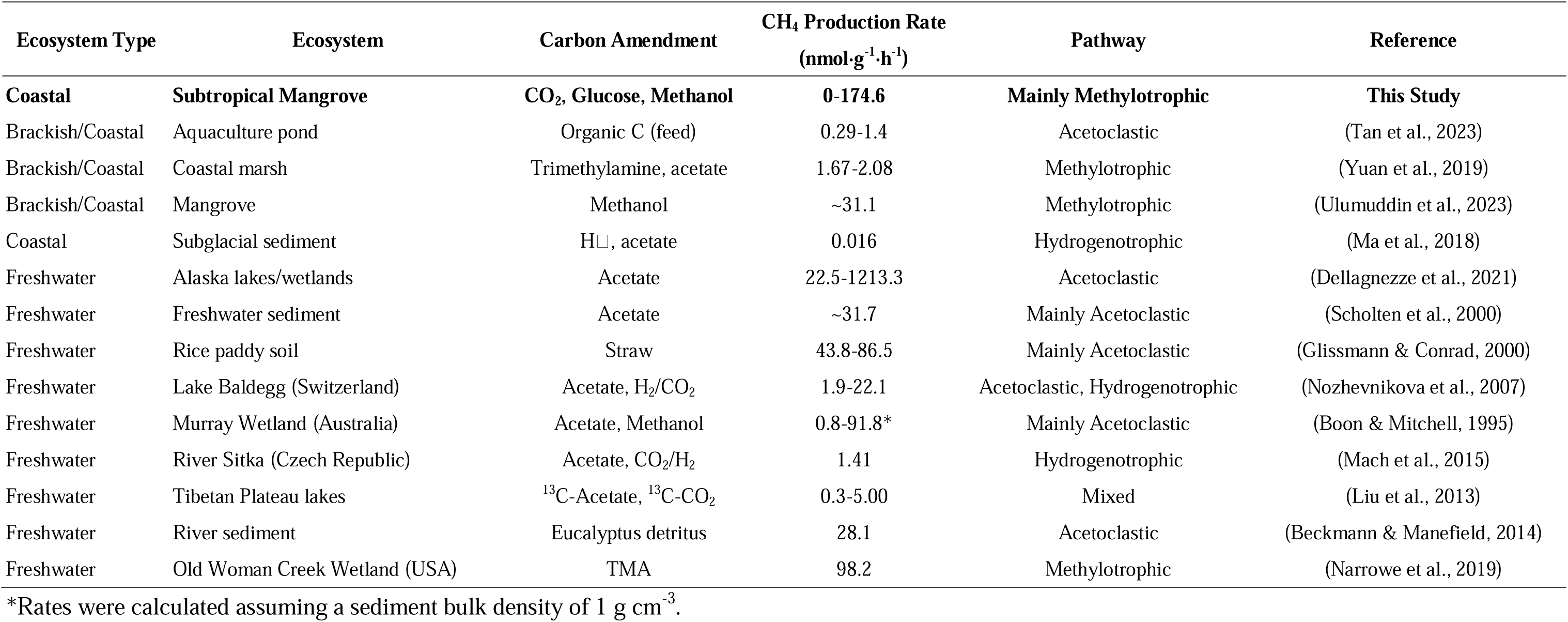
Comparisons of methane producing potential among different ecosystems.

Furthermore, the prevalence of methylotrophic and mixotrophic methanogens in coastal ecosystems has been reported in DNA-based surveys (Euler et al., 2023; Taketani et al., 2010; Yu et al., 2020; Zhou et al., 2015), supported by cultivation studies (Mohanraju et al., 1997; Zhou et al., 2023), and reflected in metatranscriptomic profiles (Cai et al., 2022; Zhang et al., 2020a; Zhang et al., 2020b),, and suggested by substrate-amended incubations (Dong et al., 2024). However, DNA- and RNA-based approaches primarily reflect genetic potential and transcriptional activity and do not directly link pathway-specific CH_4_ production with the methanogenic taxa responsible for substrate assimilation. Building on these previous observations, the present study connects pathway-specific CH_4_ production with DNA-SIP-resolved substrate-assimilating methanogenic taxa. This provides functional evidence that methylotrophic and mixotrophic methanogens, rather than acetoclastic methanogens, are potentially active contributors to CH_4_ production in mangrove soils.

This finding is consistent with a previous DNA-SIP study of methanogens using 16S metabarcoding in the coastal marsh sediments and marine sediments, which showed methylotrophic methanogens affiliated with the genus *Methanococcoides* were potentially active (Jameson et al., 2019; Yin et al., 2019). In contrast, DNA-SIP studies conducted in freshwater-dominated environments have typically reported distinct groups of potentially active methanogens, including acetoclastic *Methanosaeta* and hydrogenotrophic *Methanocella* and *Methanobacterium* in rice paddy soils (Han et al., 2018; Lee et al., 2012; Xu et al., 2017), hydrogenotrophic *Methanoregula* in freshwater wetlands (Beulig et al., 2015), and hydrogenotrophic *Methanoculleus* in bioreactor systems (Barret et al., 2015; Wang et al., 2018).

Phylogenetic analysis integrating representative *mcrA* OTUs from this study with 149 reference sequences from the NCBI database representing coastal and freshwater ecosystems showed that the potentially active methanogens identified in our ^13^C-labeled heavy fractions clustered within clades widely reported from coastal wetlands worldwide (Fig. 7a; Fig. S5). In addition, these lineages showed clear separation from those predominantly associated with terrestrial freshwater systems. Similarly, methanogenic 16S OTUs formed a distinct cluster when compared with 169 reference sequences, grouping with lineages commonly reported from coastal and saline ecosystems (Fig. 7b; Fig. S6). Based on our DNA-SIP results, we suspect that coastal ecosystems in which methylotrophic methanogens are commonly observed may harbor substantial CH_4_ production potential under saline-influenced conditions.

From an energetic perspective, methylotrophic and mixotrophic methanogenic pathways are not inherently disadvantaged under sulfate-rich conditions. Previous thermodynamic analyses indicate that methylotrophic methanogenesis can yield relatively high Gibbs free energy changes, whether via hydrogen-dependent methyl reduction (approximately −113 kJ mol^-1^) or methanol fermentation (approximately −105 kJ mol^-1^), whereas acetoclastic methanogenesis yields substantially lower energy gains (approximately −33 kJ mol^-1^) (Liu & Whitman, 2008). In comparison, SO_4_ ^2-^ reduction coupled to acetate oxidation yields Gibbs free energy changes on the order of −48 kJ mol^-1^ (Loka Bharathi, 2008). These energetic considerations suggest that methylotrophic methanogenesis can remain competitive in sulfate-rich environments where acetate-dependent pathways may be thermodynamically constrained.

In addition, metagenomic studies have identified genetic potential for lignin depolymerization (Ding et al., 2025) and O-demethylation pathways (Yu et al., 2023) in mangrove sediments, highlighting ongoing methyl-group turnover during organic matter degradation. Although such pathways do not necessarily imply direct extracellular accumulation of methanol, they indicate the potential generation of low-molecular-weight methylated intermediates that may support methylotrophic metabolisms. Consistent with this biogeochemical potential, the substantial salt-extractable methanol pool observed in the studied mangrove soils (11.3-17.2 μmol g^-1^ soil) provides additional support for the availability of endogenous methylated substrates. While this operationally defined soil pool is not directly comparable to dissolved concentrations measured in aquatic systems, it is markedly higher than methanol concentrations reported for coastal and open-ocean surface waters (0.026-0.256 nmol mL^-1^ water) (Jiang et al., 2024), suggesting that mangrove sediments may represent important reservoirs of methylated carbon relative to overlying water columns.

In addition, SRB accounted for a similar proportion of the bacterial community (approximately 2%) across the three sites in fresh soils. However, following ^13^C-based density fractionation, SRB exhibited marked enrichment in Guandu and moderate enrichment in Bali (Fig. S8), with *Desulfosporosinus* showing the most pronounced increase in Guandu relative to its corresponding fresh soil. Previous studies have shown that *Desulfosporosinus* possesses genomic features compatible with syntrophic potential (Worm et al., 2014).

Similarly, SRB within the same family (*Peptococcaceae*) have been observed to co-occur and establish syntrophic interactions with *Methanosarcina* under methanogenic conditions (Giangeri et al., 2023; Kato & Igarashi, 2019). Consistent with this ecological association, Guandu was characterized by the dominance of *Methanosarcina*, unclassified *Methanosarcinaceae*, and unclassified *Methanosarcinales* in the ^13^C-labeled fractions, whereas Bali exhibited a similar but less pronounced pattern with lower relative representation of *Methanosarcina*.

Interactions between methanogens and SRB have been reported to range from competitive to cooperative, depending on SO_4_ ^2-^ availability, substrate type, and local geochemical conditions, including potential hydrogen, formate, or mineral mediated electron transfer (Brileya et al., 2014; Giangeri et al., 2023; Großkopf et al., 2016; Kato & Igarashi, 2019). While the present study does not directly resolve the underlying mechanisms, the site-specific enrichment of SRB alongside mixotrophic methanogens suggests that interactions between methanogens and SRB may contribute to anaerobic network structure in certain estuarine mangrove soils.

In contrast, methylotrophic *Methanolobus*, which dominated the downstream sites, has rarely been implicated in direct syntrophic associations with SRB. Together, these observations suggest that SRB enrichment in the ^13^C fractions may reflect site-specific anaerobic network configurations rather than uniform competitive exclusion. Future studies may consider focusing on the coupling between SRB and methanogens in mangrove ecosystems.

### 4.3 Decoupling of CH_4_ production and emission in saline influenced mangrove ecosystems

Field-based observations and global syntheses have consistently demonstrated that saline coastal wetlands exhibit substantially lower surface CH_4_ emissions compared to freshwater ecosystems (Al-Haj & Fulweiler, 2020; Poffenbarger et al., 2011; Shiau & Chiu, 2020). This apparent discrepancy between low net CH_4_ fluxes and the substantial methanogenic potential observed in mangrove soils of this present study underscores the limitation of interpreting surface efflux measurements in isolation.

Our previous studies of the aerobic methanotrophic communities (i.e. methane oxidizing bacteria, MOB) in the same mangrove forests (Shiau et al., 2020; Shiau et al., 2017a), together with a coastline-scale survey of mangrove forests spanning 370 km of Taiwan (Shiau et al., 2025), consistently identified potentially active MOB dominated by members of the family *Methylomonadaceae* (i.e. Type Ia methanotrophs) across a wide range of environmental conditions. Type Ia methanotrophs have been suggested to exhibit traits consistent with r-strategist life-history strategies and are often associated with environments characterized by elevated CH_4_ availability (Ho et al., 2013; Knief, 2015). Furthermore, a growing body of DNA-SIP studies targeting MOB in saline environments worldwide, such as mangroves, salt marshes, estuarine sediments, and saline lake soils, indicates that Type Ia methanotrophs represent a common and potentially active component of CH_4_ oxidation under saline conditions (Table S3) (Deng et al., 2019; do Carmo Linhares et al., 2021; Fang et al., 2022; Fang et al., 2025; Shiau et al., 2025; Shiau et al., 2020; Shiau et al., 2017a).

When considered together with our previous SIP-based investigations of MOB in the same mangrove ecosystems at Guandu, Zhuwei, and Bali under comparable temperature conditions (25 °C) (Shiau et al., 2020; Shiau et al., 2017a), which reported CH_4_ oxidation potentials of 0.010, 0.026, and 0.052 μmol g^-1^ h^-1^ for these respective sites, our results indicate that CH_4_ oxidation and production processes may occur at similar orders of magnitude across the same environmental gradient. These findings suggest that mangrove soils may function as systems characterized by rapid internal CH_4_ cycling, in which low net CH_4_ efflux reflects the balance between production and consumption processes rather than intrinsically low methanogenic activity. This suggestion is also in line with recently published studies using δ^13^C-CH_4_, showing the importance of CH_4_ oxidation in reducing the net CH_4_ effluxes from mangrove ecosystems (Cotovicz et al., 2024; Euler et al., 2023).

Furthermore, several studies have suggested that Type I methanotrophs are generally psychrophilic and community composition shifting under altered thermal regimes (Chen & Shiau, 2025; Roldán et al., 2022; Urmann et al., 2009). This raises the possibility that the balance between CH_4_ production and oxidation in mangrove soils could shift under future climate warming. Future research should carefully evaluate how this CH_4_ cycling balance responds to rising temperatures.

In addition to SRB, other microbial groups may also influence the availability of methylated substrates in mangrove soils. In the present study, *Methyloceanibacter* and other methylotrophs were consistently detected in the ^13^C-labeled heavy fractions, suggesting potentially active incorporation of C1 compounds under saline environments (Fig. S9). Members of the genus *Methyloceanibacter* have been reported to utilize methanol and other methylated substrates (Takeuchi et al., 2014; Takeuchi et al., 2019; Vekeman et al., 2016), raising the possibility that competition for methanol may also occur between methylotrophic bacteria and methanogens in certain saline environments. Although the relative importance of such interactions remains unclear, this observation highlights the potential complexity of methylated carbon partitioning within saline-influenced mangrove soils.

In addition, anaerobic methane-oxidizing archaea (ANME) and other taxa associated with anaerobic oxidation of CH_4_ (AOM) have been reported as important CH_4_ consumers in certain sulfate-rich saline environments (Bhattarai et al., 2017; Wallenius et al., 2021; Wang et al., 2019). In the present study, however, AOM-related 16S rRNA gene sequences were detected only at very low abundances in both fresh soils and SIP incubations (i.e. <0.07% of the relative abundance). This pattern may reflect the experimental design, as density fractionation was guided by *mcrA*-based qPCR targeting methanogenic populations. In addition, the low representation of AOM-related taxa may also reflect ecological constraints, as ANME are generally slow-growing microorganisms that may be less competitive in highly dynamic, tidally influenced environments (Wallenius et al., 2021). Thus, the contribution of AOM-mediated CH_4_ oxidation was not explicitly incorporated into the mechanistic framework proposed in this study. In high-turnover CH_4_ systems such as mangrove soils, the potential contribution of AOM remains an open question and warrants further investigation.

Overall, these findings indicate that CH_4_ production in saline mangrove blue carbon ecosystems can be sustained by methylotrophic and mixotrophic methanogenic pathways while coexisting with efficient CH_4_ consumption. This pattern suggests that low net CH_4_ emissions may not necessarily reflect weak methanogenesis, but may instead arise from coupled CH_4_ production and consumption within mangrove soils. By linking pathway-specific CH_4_ production with DNA-SIP-resolved substrate-assimilating methanogens, this study provides a mechanistic basis for interpreting low surface CH_4_ emissions in saline mangrove ecosystems. Future *in situ* isotope probing or ^13^C tracer experiments under natural conditions will help bridge these incubation-based findings to ecosystem-scale CH_4_ fluxes. Clarifying how warming alters the balance between CH_4_ production and consumption will further improve predictions of mangrove CH_4_ emissions under future climate scenarios.

## Acknowledgements

This study is funded by the National Science and Technology Council, Taiwan [110-2313-B-002-033-MY3, 113-2313-B-002-031-MY3 and 113-2321-B-002-037] and Fisheries Agency, Ministry of Agriculture, Taiwan [115AS-13.3.1-FA-01]. The authors thank Yi-Hsuan Huang, Pei-Yi Yu and Ting-Kai Chen for helping with the field sampling.

## Author Contributions

Y.-W. Z.: Investigation, Methodology, Formal analysis, Data curation, Visualization, Writing – original draft. Y.-J. S.: Conceptualization, Methodology, Formal analysis, Writing – review and editing, Supervision, Funding acquisition, Validation. Y.-W. Z. and Y.-J. S. conducted the field experiments and interpreted the data. All authors have read and approved the final version of the manuscript.

## Declaration of generative AI and AI-assisted technologies in the manuscript preparation process

During the preparation of this manuscript, the authors used ChatGPT 5.5 solely for language refinement. All scientific content, data interpretation, wording choices, and conclusions were reviewed, edited, and approved by the authors, who take full responsibility for the final version of the manuscript.

