## Supplemental Information for "DNA-SIP reveals salinity-associated niche differentiation of potentially active methanogens in mangrove soils"

**This Supporting Information file includes:**

- **Tables:** 3 (Tables S1–S3)
- **Figures:** 9 (Figures S1–S9)

**Table S1** Differential abundance results for the 50 most abundant *mcrA* OTUs in the field mangrove soils using ALDEx2. Statistically significant differences (FDR-adjusted $p < 0.05$) and biologically meaningful effect sizes (|effect size| > 1) are summarized as pairwise site comparisons. Blank cells indicate no statistically significant or biologically meaningful difference was detected.

| Taxon | Statistical Significance ($P < 0.05$) | Biological Effect (\|Effect size\|>1) |
|---|---|---|
| OTU_001_*Methanolobus*_unclassified | Zhuwei > Guandu | Zhuwei = Bali > Guandu |
| OTU_002_*Methanolobus_vulcani* | | |
| OTU_003_uncultured_*Methanosarcinales*_archaeon_unclassified | | Guandu = Bali > Zhuwei |
| OTU_004_*Methanolobus_profundi* | | Zhuwei > Bali > Guandu |
| OTU_005_*Methanolobus*_unclassified | | Zhuwei > Guandu |
| OTU_006_*Methanolobus*_unclassified | Zhuwei > Guandu | Zhuwei = Bali > Guandu |
| OTU_007_*Methanolobus*_unclassified | Zhuwei > Guandu | Zhuwei > Bali = Guandu |
| OTU_008_*Methanolobus*_unclassified | | Bali > Guandu |
| OTU_009_*Methanosarcina_acetivorans* | | Guandu > Zhuwei |
| OTU_010_*Methanosarcina_acetivorans* | | |
| OTU_011_Archaea_unclassified | | |
| OTU_012_*Methanolobus*_profundi | | |
| OTU_013_*Methanolobus*_unclassified | Zhuwei > Guandu = Bali | Zhuwei > Guandu = Bali |
| OTU_014_*Methanosarcinaceae*_unclassified | | |
| OTU_015_*Methanolobus_profundi* | Zhuwei > Guandu | Zhuwei > Guandu |
| OTU_016_*Methanosarcinaceae*_unclassified | | |
| OTU_017_*Methanosarcina_barkeri* | | |
| OTU_018_*Methanolobus*_unclassified | | |
| OTU_019_*Methanomicrobiaceae*_unclassified | | |
| OTU_020_*Methanosarcina*_unclassified | | |
| OTU_021_*Methanosarcinaceae*_unclassified | | Guandu > Bali |
| OTU_022_archaeon_enrichment_culture_clone_McrA2_unclassified | | Bali > Zhuwei = Guandu |
| OTU_023_*Methanofollis_ethanolicus* | | |
| OTU_024_*Methanococcoides*_unclassified | | Bali > Guandu |
| OTU_025_*Methanolobus*_unclassified | | |
| OTU_026_*Methanosarcinales*_unclassified | | Guandu > Zhuwei = Bali |
| OTU_027_uncultured_archaeon | | |
| OTU_028_*Methanosarcina*_unclassified | | |
| OTU_029_*Methanolobus*_unclassified | | Zhuwei = Bali > Guandu |
| OTU_030_*Methanofollis_ethanolicus* | | |
| OTU_031_*Methanolobus*_unclassified | Zhuwei > Guandu | Zhuwei = Bali > Guandu |
| OTU_032_*Methanosarcinaceae*_unclassified | | Bali > Guandu |
| OTU_033_*Methanolobus*_unclassified | Zhuwei > Guandu | Zhuwei > Bali = Guandu |
| OTU_034_*Methanomicrobiales*_unclassified | | |
| OTU_035_uncultured_*Methanosarcinales*_archaeon_unclassified | | |
| OTU_036_*Methanolobus_profundi* | | |
| OTU_037_*Methanosarcinaceae*_unclassified | | Guandu > Bali |
| OTU_038_*Methanosarcinaceae*_unclassified | | |
| OTU_039_*Methanofollis_ethanolicus* | | |
| OTU_040_*Methanosarcinaceae*_unclassified | | |
| OTU_041_*Methanosarcinaceae*_unclassified | | |
| OTU_042_*Methanolobus*_unclassified | | |
| OTU_043_*Methanosarcinaceae*_unclassified | | |
| OTU_044_*Methanomicrobiaceae*_unclassified | | |
| OTU_045_Archaea_unclassified | | |
| OTU_046_Archaea_unclassified | | |
| OTU_047_Archaea_unclassified | | |
| OTU_048_*Methanosarcinaceae*_unclassified | | Guandu > Bali |
| OTU_049_*Methanolobus*_unclassified | | |
| OTU_050_*Methanomicrobiales*_unclassified | | |

**Table S2** Differential abundance results for the 50 most abundant *mcrA* OTUs in the $^{13}$C-methanol labeled DNA heavy fractions using ALDEx2. Statistically significant differences (FDR-adjusted p < 0.05) and biologically meaningful effect sizes (|effect size| > 1) are summarized as pairwise site comparisons. Blank cells indicate no statistically significant or biologically meaningful difference was detected.

| Taxon | Statistical Significance (*P* < 0.05) | Biological Effect (\|Effect size\|>1) |
|---|---|---|
| OTU_001_*Methanolobus*_unclassified | | |
| OTU_002_*Methanolobus_vulcani* | | Guandu > Zhuwei = Bali |
| OTU_003_uncultured_*Methanosarcinales*_archaeon_unclassified | | Guandu > Zhuwei |
| OTU_004_*Methanolobus_profundi* | Zhuwei > Guandu | Zhuwei > Bali > Guandu |
| OTU_005_*Methanolobus*_unclassified | | Zhuwei = Bali > Guandu |
| OTU_006_*Methanolobus*_unclassified | | Zhuwei > Guandu |
| OTU_007_*Methanolobus*_unclassified | Zhuwei > Guandu | Zhuwei = Bali > Guandu |
| OTU_008_*Methanolobus*_unclassified | | |
| OTU_009_*Methanosarcina_acetivorans* | | Guandu > Zhuwei = Bali |
| OTU_010_*Methanosarcina_acetivorans* | | |
| OTU_011_Archaea_unclassified | | Guandu > Zhuwei |
| OTU_012_*Methanolobus*_profundi | | Bali > Guandu |
| OTU_013_*Methanolobus*_unclassified | | Zhuwei > Bali > Guandu |
| OTU_014_*Methanosarcinaceae*_unclassified | | Guandu > Zhuwei |
| OTU_015_*Methanolobus_profundi* | | |
| OTU_016_*Methanosarcinaceae*_unclassified | | |
| OTU_017_*Methanosarcina_barkeri* | | |
| OTU_018_*Methanolobus*_unclassified | | Guandu > Zhuwei = Bali |
| OTU_019_*Methanomicrobiaceae*_unclassified | | |
| OTU_020_*Methanosarcina*_unclassified | | |
| OTU_021_*Methanosarcinaceae*_unclassified | | |
| OTU_022_archaeon_enrichment_culture_clone_McrA2_unclassified | | Bali > Guandu |
| OTU_023_*Methanofollis_ethanolicus* | | |
| OTU_024_*Methanococcoides*_unclassified | | |
| OTU_025_*Methanolobus*_unclassified | | |
| OTU_026_*Methanosarcinales*_unclassified | | |
| OTU_027_uncultured_archaeon | | |
| OTU_028_*Methanosarcina*_unclassified | | Guandu > Zhuwei = Bali |
| OTU_029_*Methanolobus*_unclassified | | Zhuwei > Guandu |
| OTU_030_*Methanofollis_ethanolicus* | | |
| OTU_031_*Methanolobus*_unclassified | | Zhuwei > Guandu |
| OTU_032_*Methanosarcinaceae*_unclassified | | |
| OTU_033_*Methanolobus*_unclassified | | Zhuwei > Guandu = Bali |
| OTU_034_*Methanomicrobiales*_unclassified | | |
| OTU_035_uncultured_*Methanosarcinales*_archaeon_unclassified | | |
| OTU_036_*Methanolobus_profundi* | | |
| OTU_037_*Methanosarcinaceae*_unclassified | | |
| OTU_038_*Methanosarcinaceae*_unclassified | | |
| OTU_039_*Methanofollis_ethanolicus* | | |
| OTU_040_*Methanosarcinaceae*_unclassified | | |
| OTU_041_*Methanosarcinaceae*_unclassified | | Guandu > Zhuwei |
| OTU_042_*Methanolobus*_unclassified | | |
| OTU_043_*Methanosarcinaceae*_unclassified | | |
| OTU_044_*Methanomicrobiaceae*_unclassified | | |
| OTU_045_Archaea_unclassified | | |
| OTU_046_Archaea_unclassified | | |
| OTU_047_Archaea_unclassified | | |
| OTU_048_*Methanosarcinaceae*_unclassified | | |
| OTU_049_*Methanolobus*_unclassified | | |
| OTU_050_*Methanomicrobiales*_unclassified | | |

**Table S3** Comparisons of potentially active methanotrophs across different ecosystems using DNA stable isotope probing

| Ecosystem* | Type | Location | Major Potentially Active Methanotrophs (DNA-SIP based) | Methane oxidation potential (μmol $g^{-1}$ $d^{-1}$) | Reference |
|---|---|---|---|---|---|
| Lakeshore wetland | Saline | Qinghai Lake, China | **Type Ia:** *Methylomicrobium*, *Methylovulum* | | (Fang et al., 2022) |
| Coastal mangrove forest | Saline | Tamsui Estuary (Zhuwei), Taiwan | **Type Ia:** *Methylobacter*, *Methylosarcina*, *Methylomonas* **Type Ib:** deep-sea-5 cluster | 0.60-0.94 | (Shiau et al., 2017) |
| Riparian & Lakeshore | Saline | Yangtze River Basin, China | Riparian (Saline-adapted): **Type Ia:** *Methylobacter* | | (Fang et al., 2025) |
| Mangrove | Saline | Bertioga, São Paulo, Brazil | **Type Ia:** *Methylomonas*, *Methylobacter*, *Methylomicrobium* | 35 | (do Carmo Linhares et al., 2021) |
| Estuarine Mangrove forest | Saline | Tamsui Estuary (Guandu & Bali), Taiwan | **Type Ia:** *Methylobacter*, *Methylomonas*, unclassified *Methylococcaceae*, *Methylomicrobium* | 0.24-1.44 | (Shiau et al., 2020) |
| Tidal marsh | Saline | Yancheng, Jiangsu, China | **Type Ia:** Unclassified *Methylococcaceae*, *Methylomonas*, *Methylohalobius* | Mudflat: 2.67 Vegetated: 1.6–2.18 | (Deng et al., 2019) |
| Coastal Mangrove forest | Saline | West coast of Taiwan | **Type Ia:** Unclassified *Methylomonadaceae*, *Methylobacter*, *Methylomonas*, *Methyloceanibacter* | 0.24-1.44 | (Shiau et al., 2025) |

* Ecosystem descriptions were retained as originally reported in the cited studies.

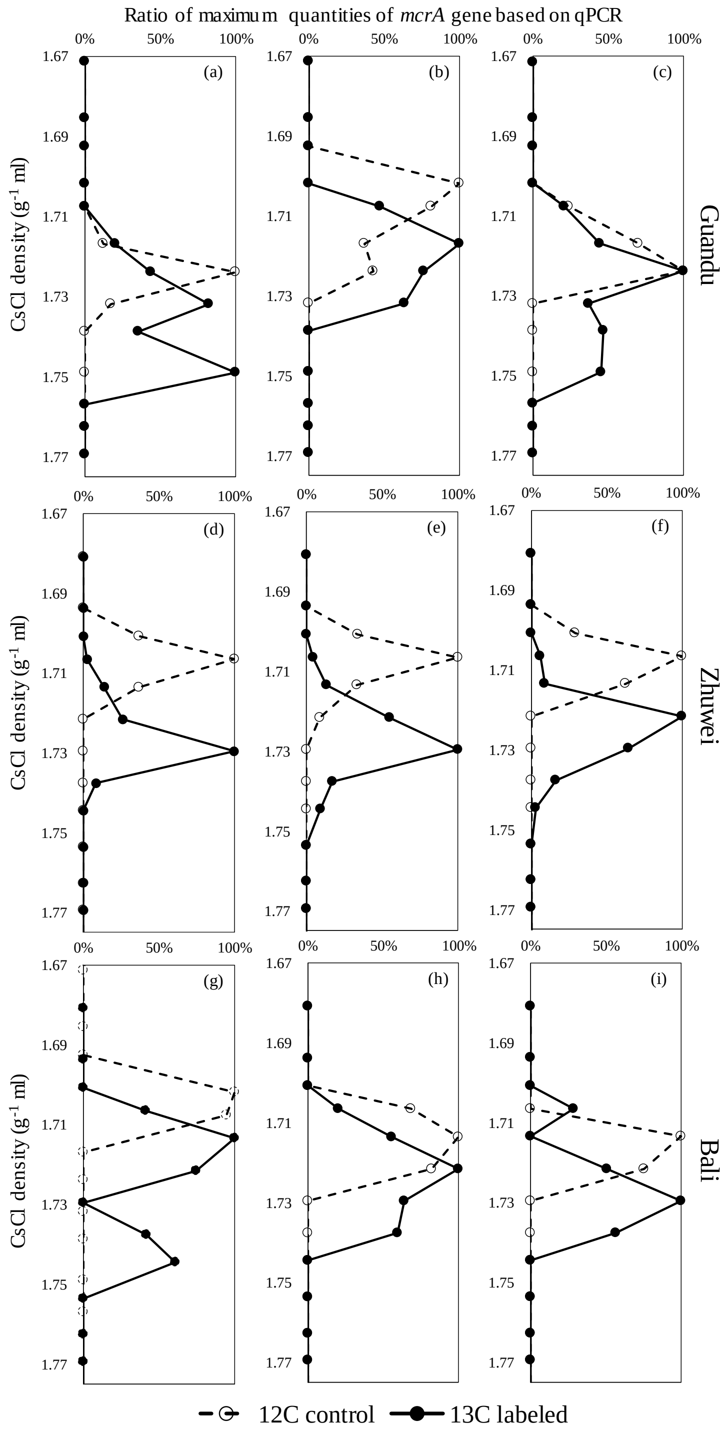


**Figure S1** Relative *mcrA* gene abundance across CsCl buoyant density fractions in $^{12}C$ control and $^{13}C$-labeled mangrove soils. For each sample, *mcrA* copy numbers in individual density fractions are expressed as a percentage of the maximum copy number detected within that sample.

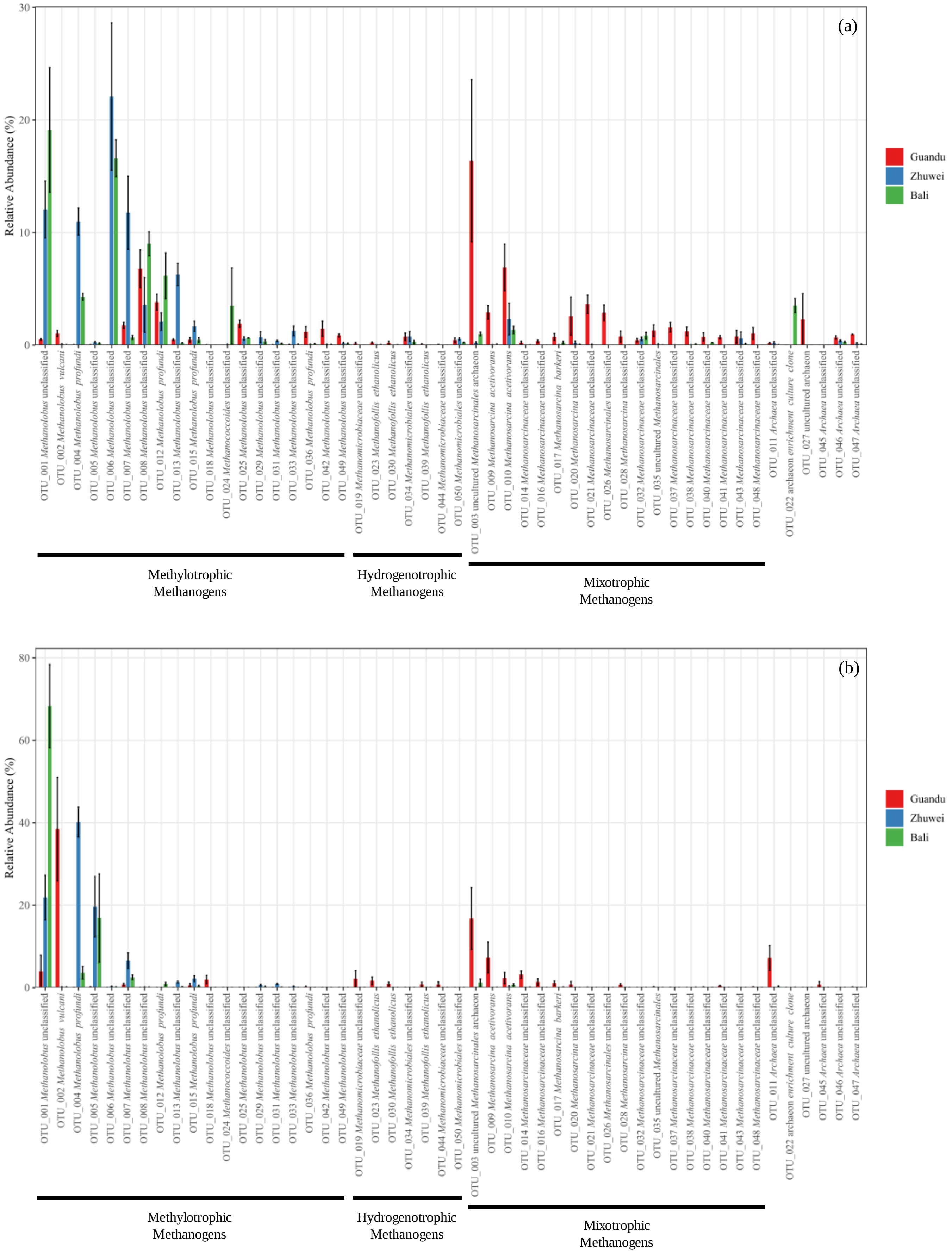


**Figure S2** Relative abundance of the 50 most abundant *mcrA* OTUs across the three mangrove forests. (a) Field soil communities. (b) $^{13}C$-labeled heavy DNA fractions obtained from DNA-SIP incubations.

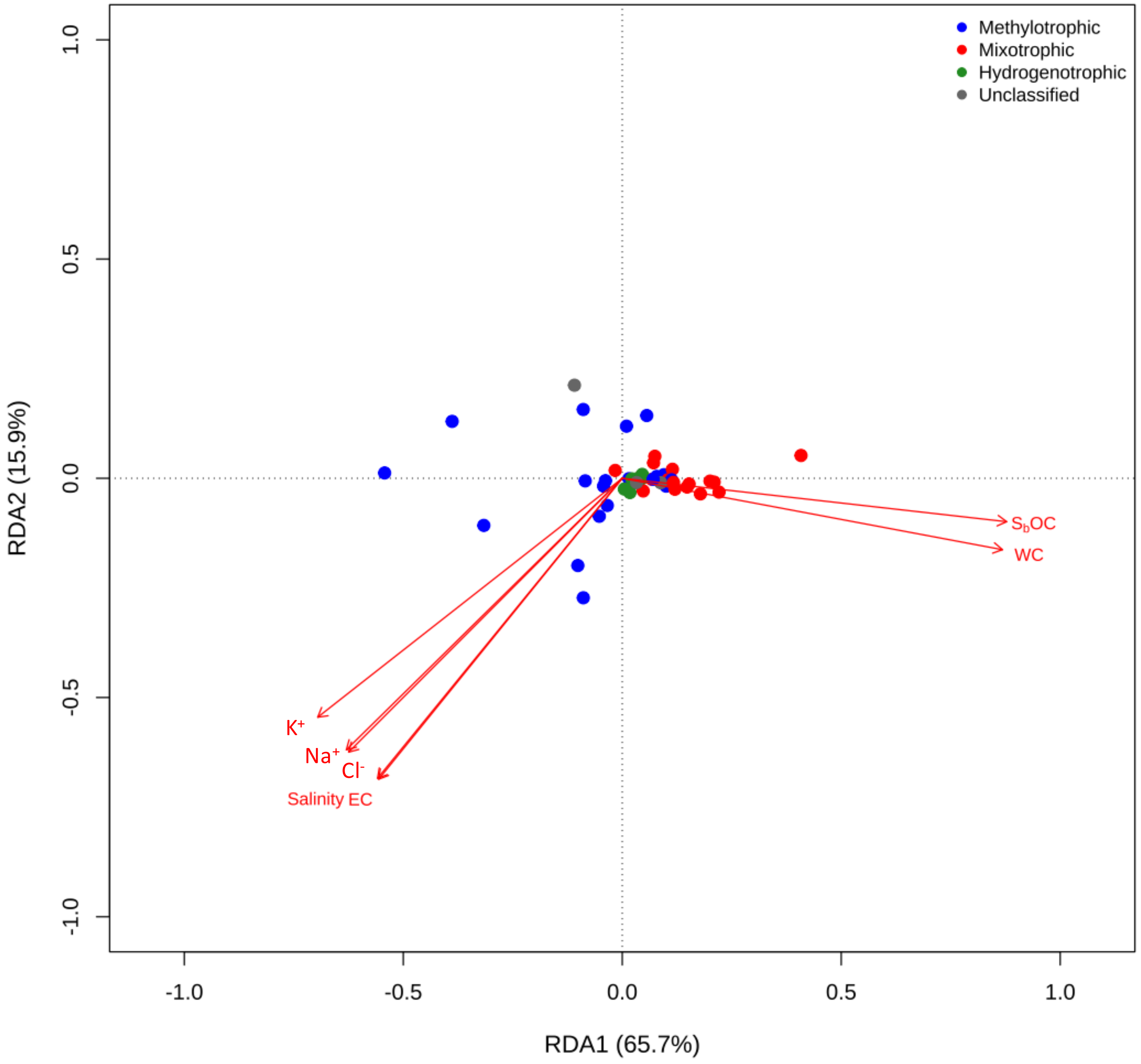


**Figure S3** Redundancy analysis (RDA) showing the relationships between methanogenic OTUs based on *mcrA* gene sequences and soil physicochemical properties in field soils from the three mangrove forests. Methylotrophic methanogens are shown in blue, mixotrophic methanogens in red, hydrogenotrophic methanogens in green, and unclassified methanogens in grey.

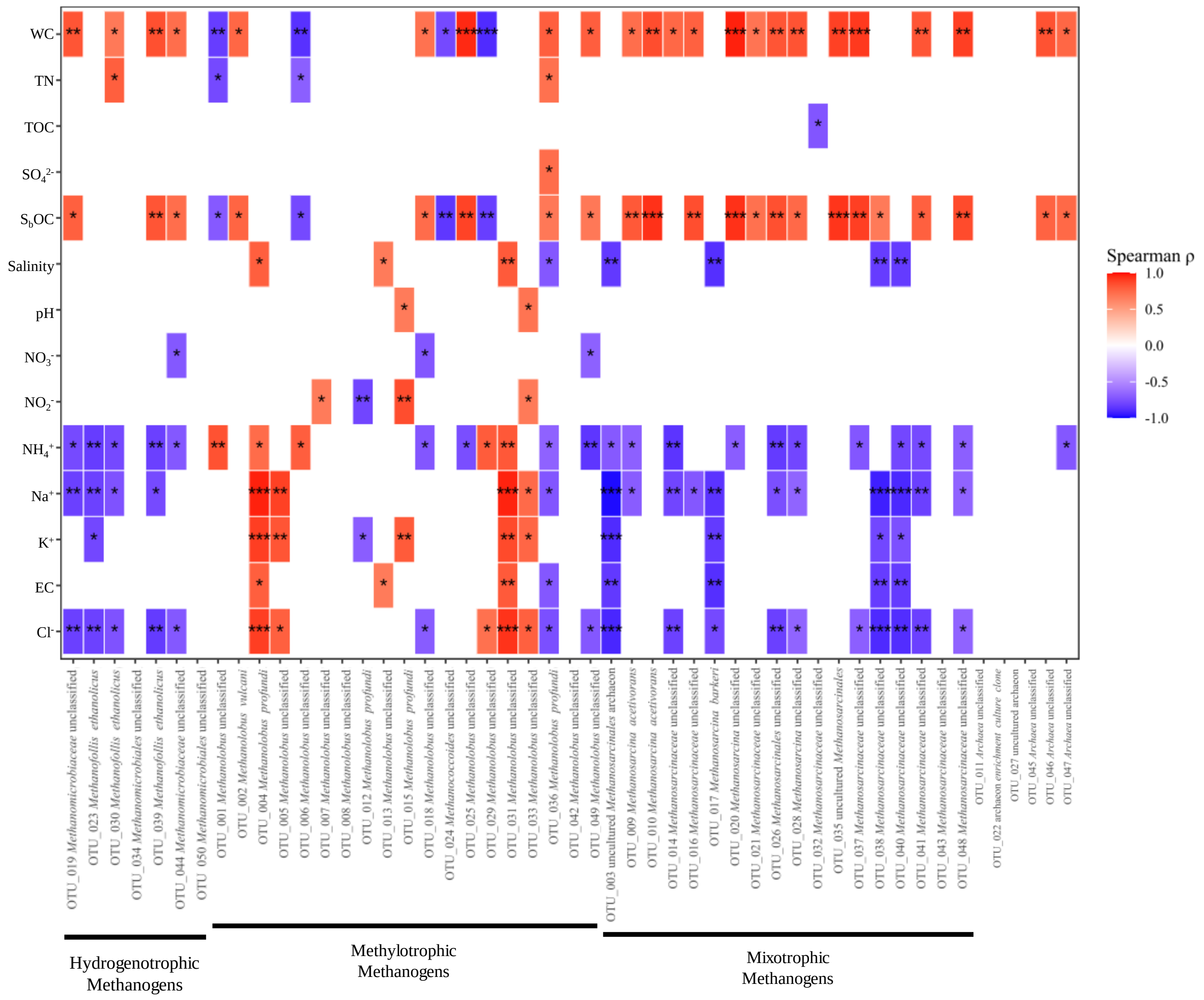


**Figure S4** Spearman correlation analysis between soil physicochemical properties and methanogenic OTUs based on *mcrA* gene sequence in field soils from three mangrove forests along the estuarine gradient in northern Taiwan.

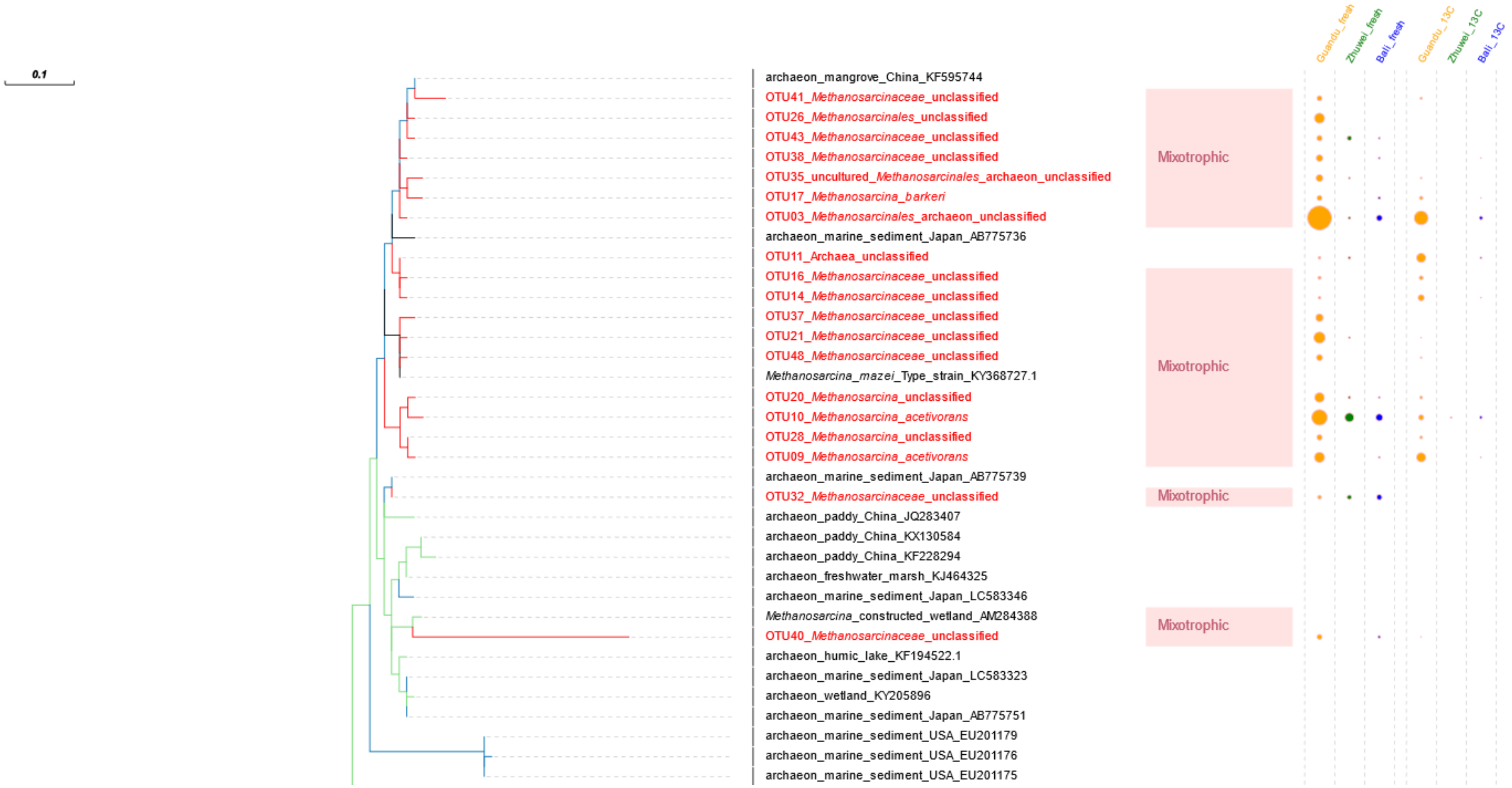


**Figure S5** Phylogenetic tree of methanogenic archaea inferred from *mcrA* gene sequences. The tree includes *mcrA* sequences recovered from mangrove forest soils in Taiwan in this study together with 149 reference sequences retrieved from the NCBI (GenBank) database, representing methanogens from freshwater and saline environments across diverse geographic regions worldwide. Amino acid sequences were aligned, and phylogenetic reconstruction was performed using the maximum-likelihood method under the WAG substitution model.

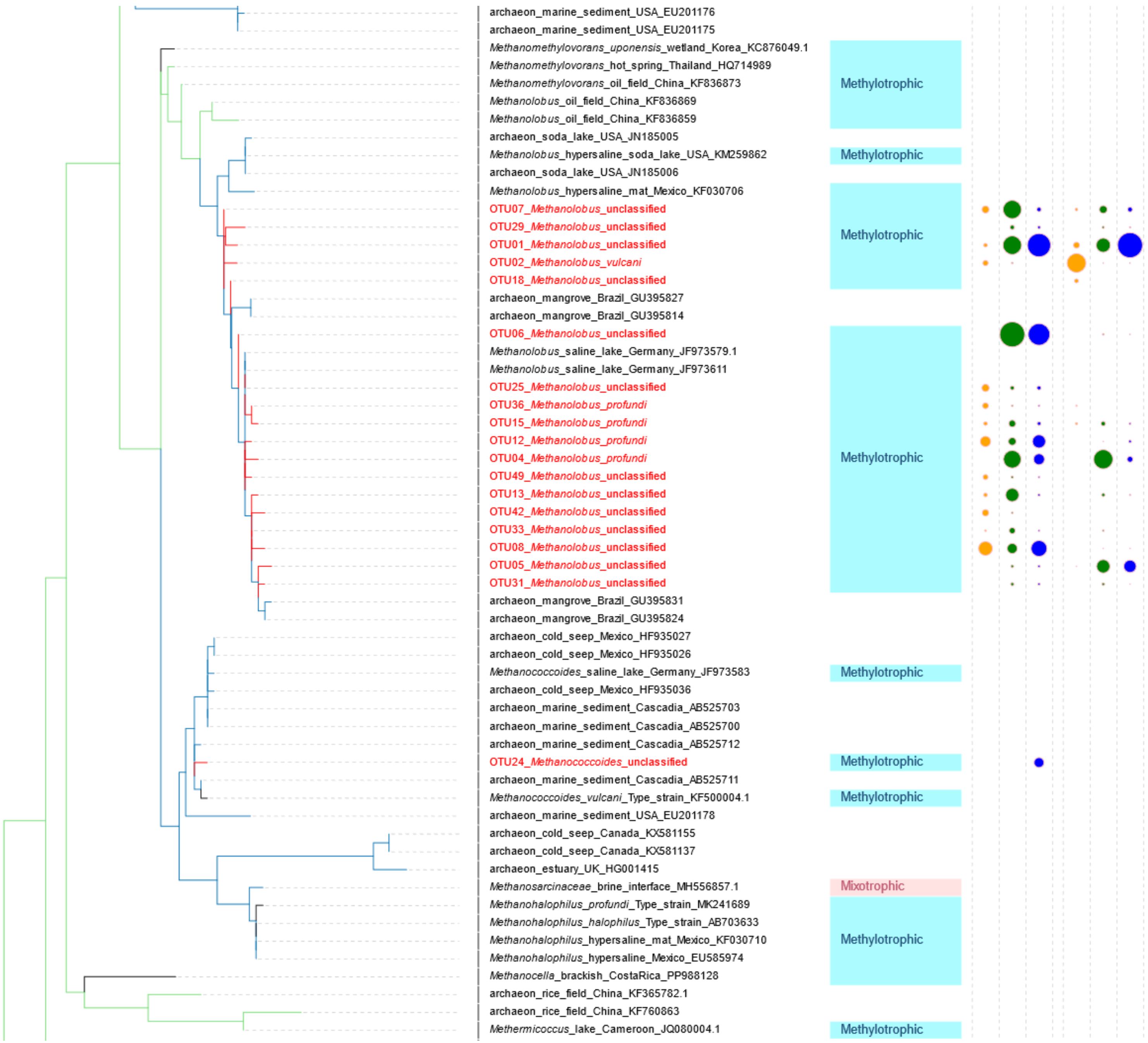


**Figure S5** continued

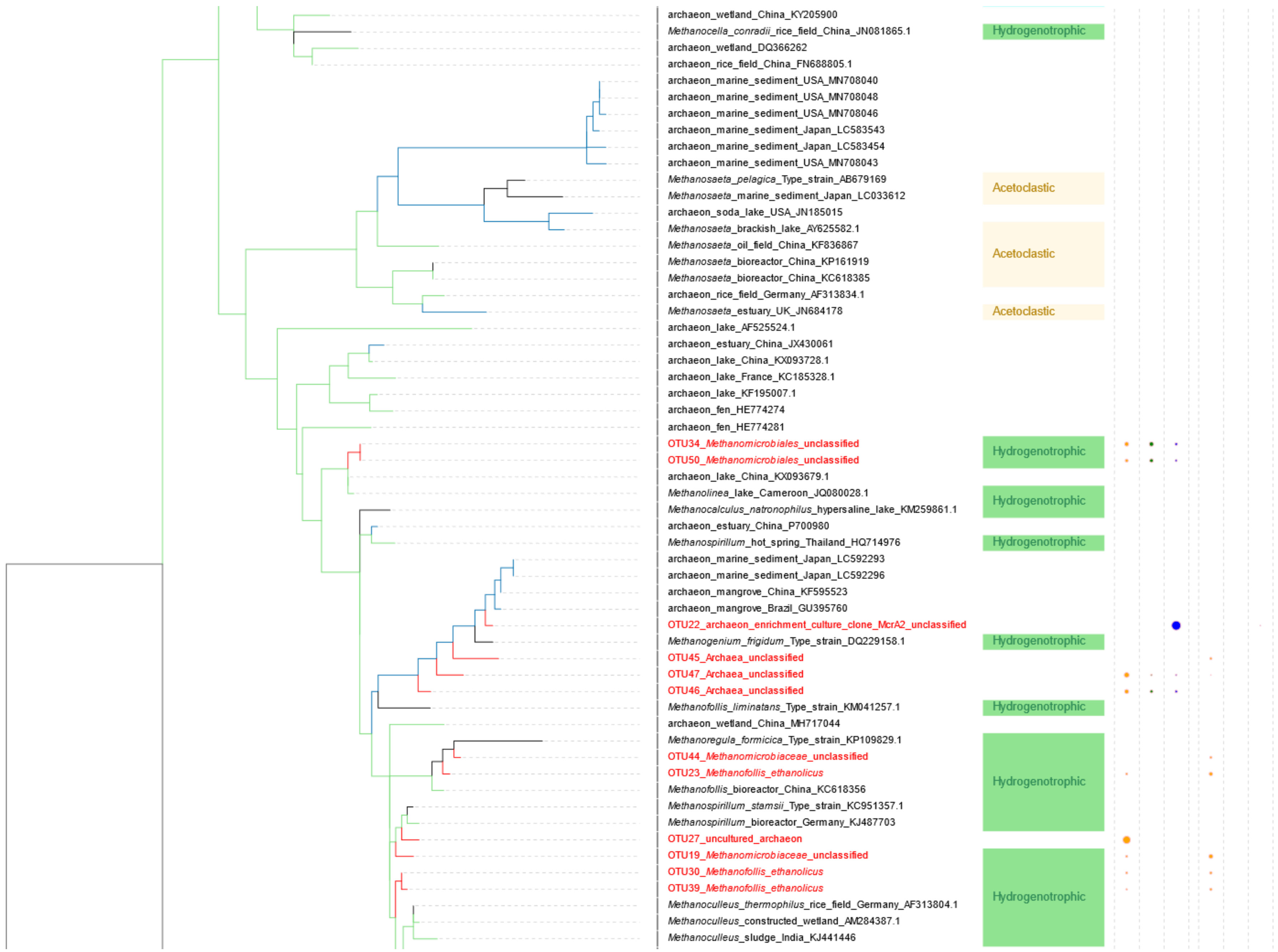


**Figure S5** continued

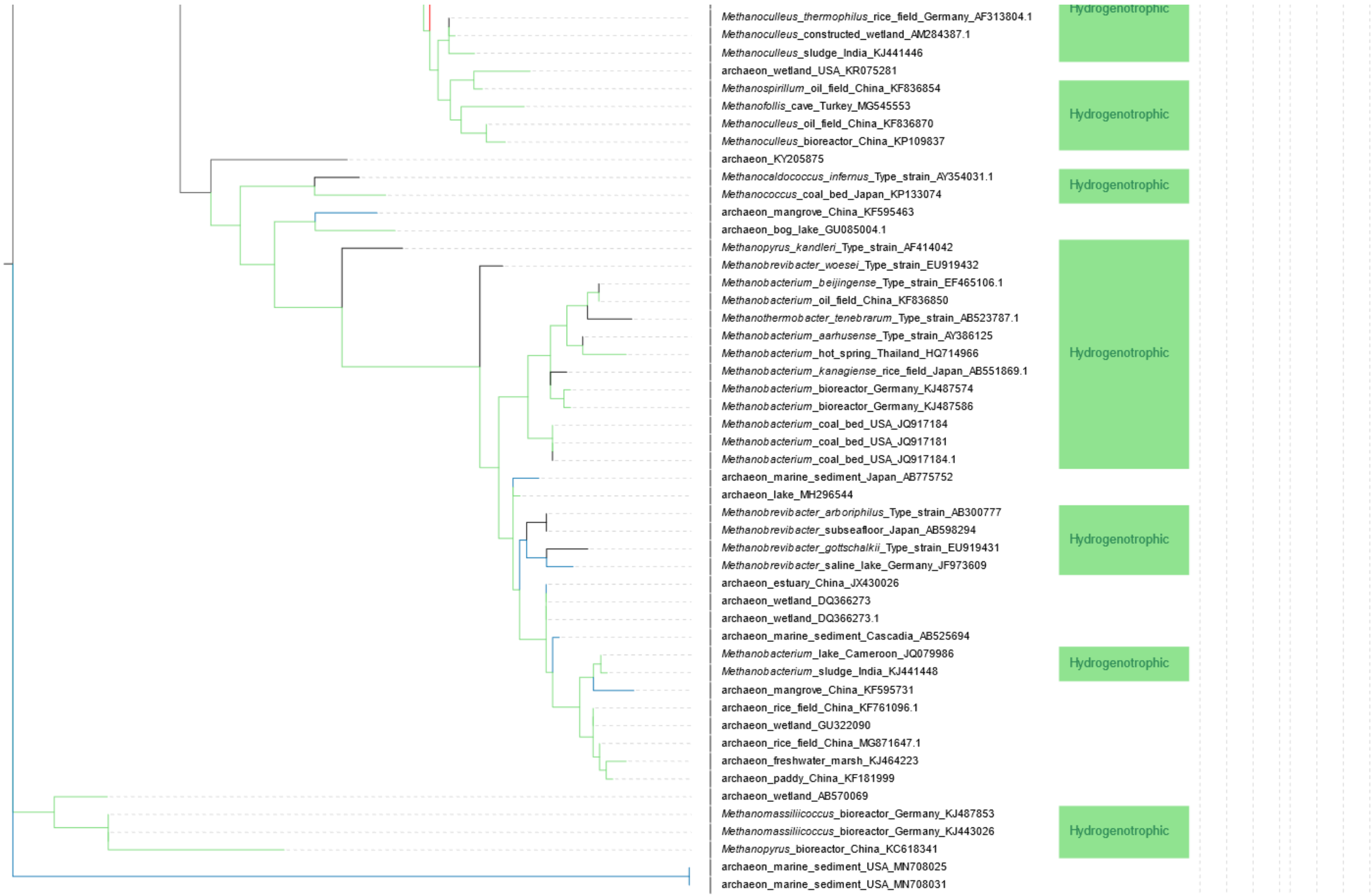


**Figure S5** continued

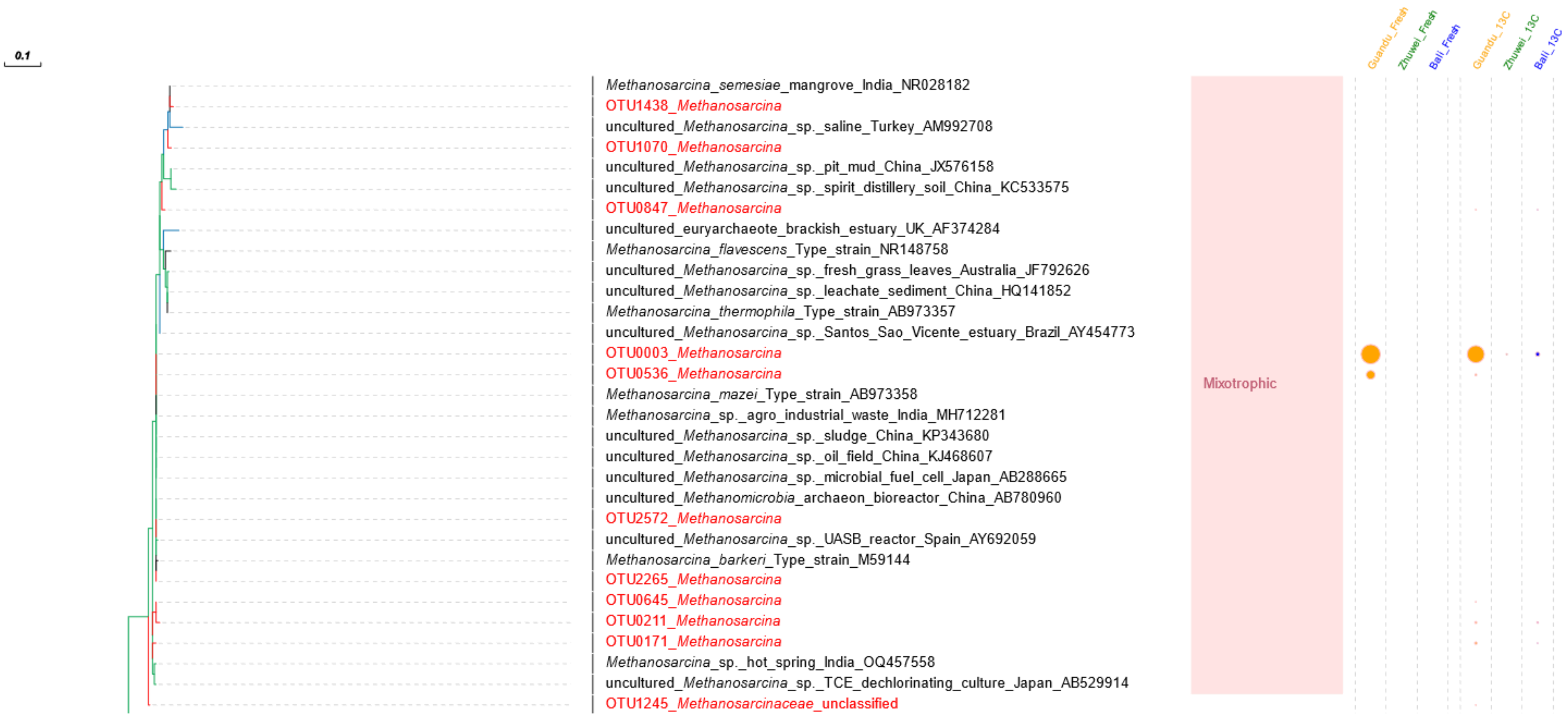


**Figure S6** Phylogenetic tree of methanogenic archaea inferred from 16S rRNA gene sequences. The tree includes the 55 most abundant methanogenic 16S rRNA gene OTUs recovered from mangrove forest soils in Taiwan in this study together with 169 reference sequences retrieved from the NCBI (GenBank) database, representing methanogens from freshwater and saline environments across diverse geographic regions worldwide. Phylogenetic reconstruction was performed using the maximum-likelihood method under the Tamura–Nei substitution model.

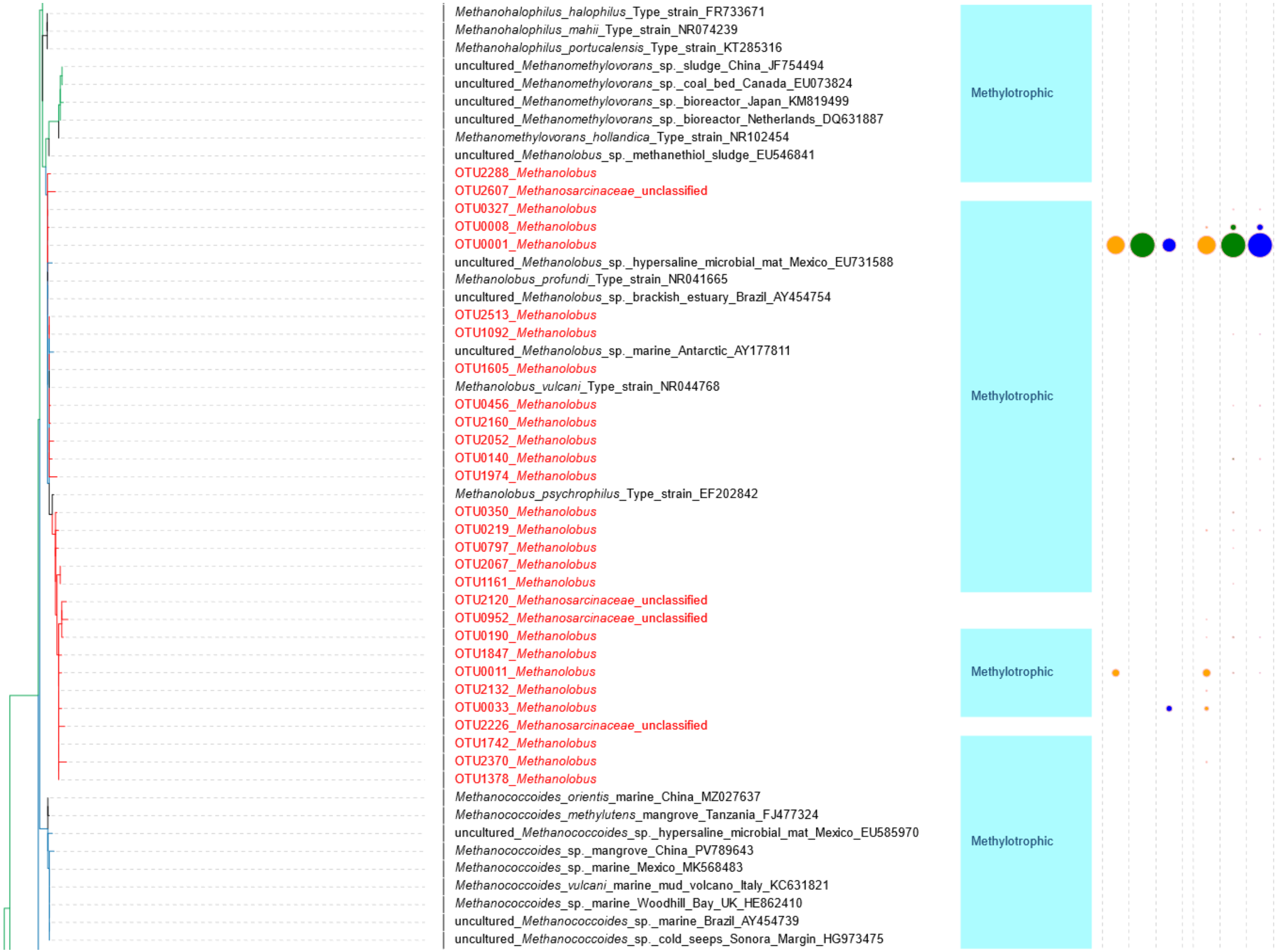


**Figure S6** continued

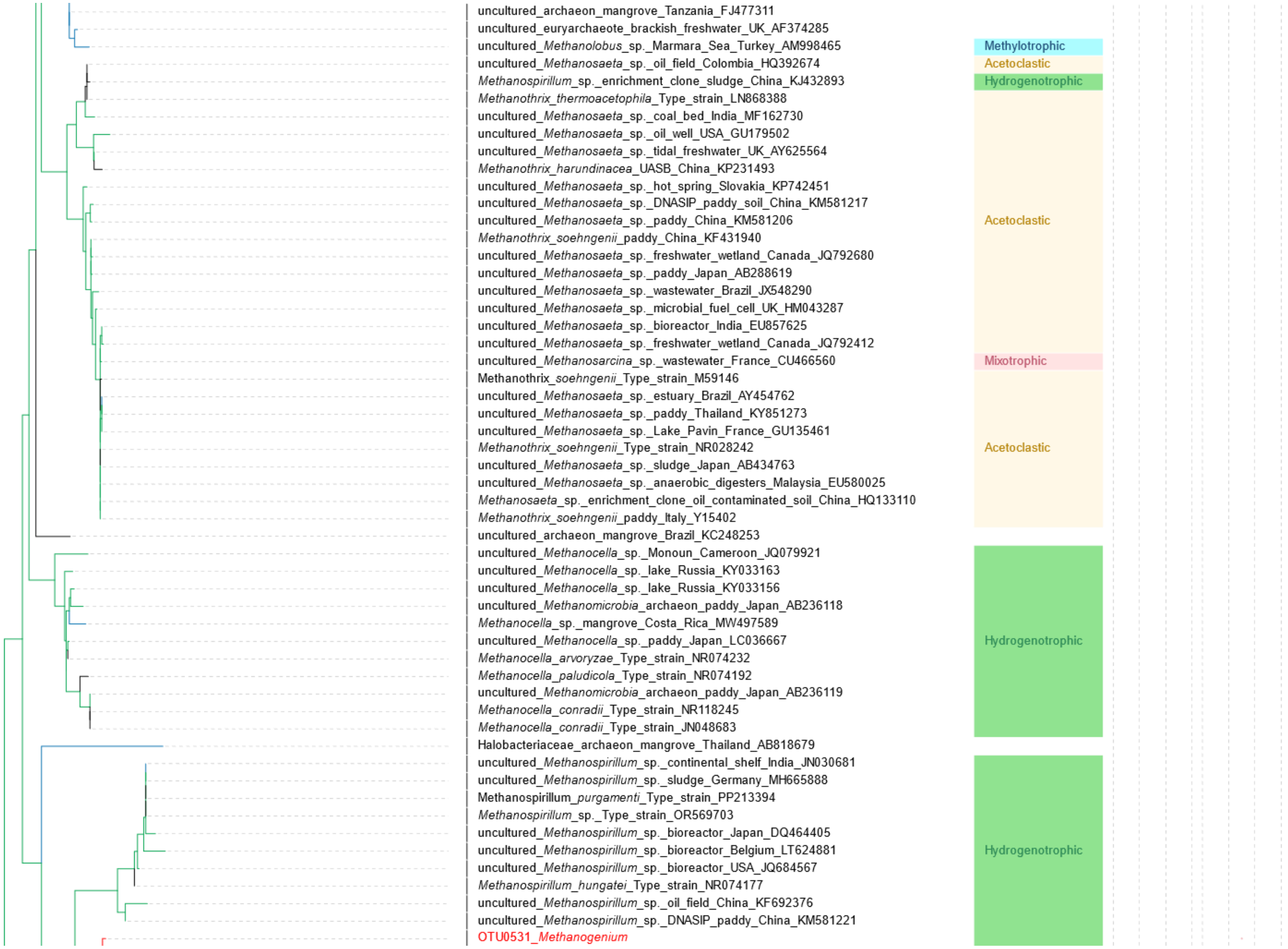


**Figure S6** continued

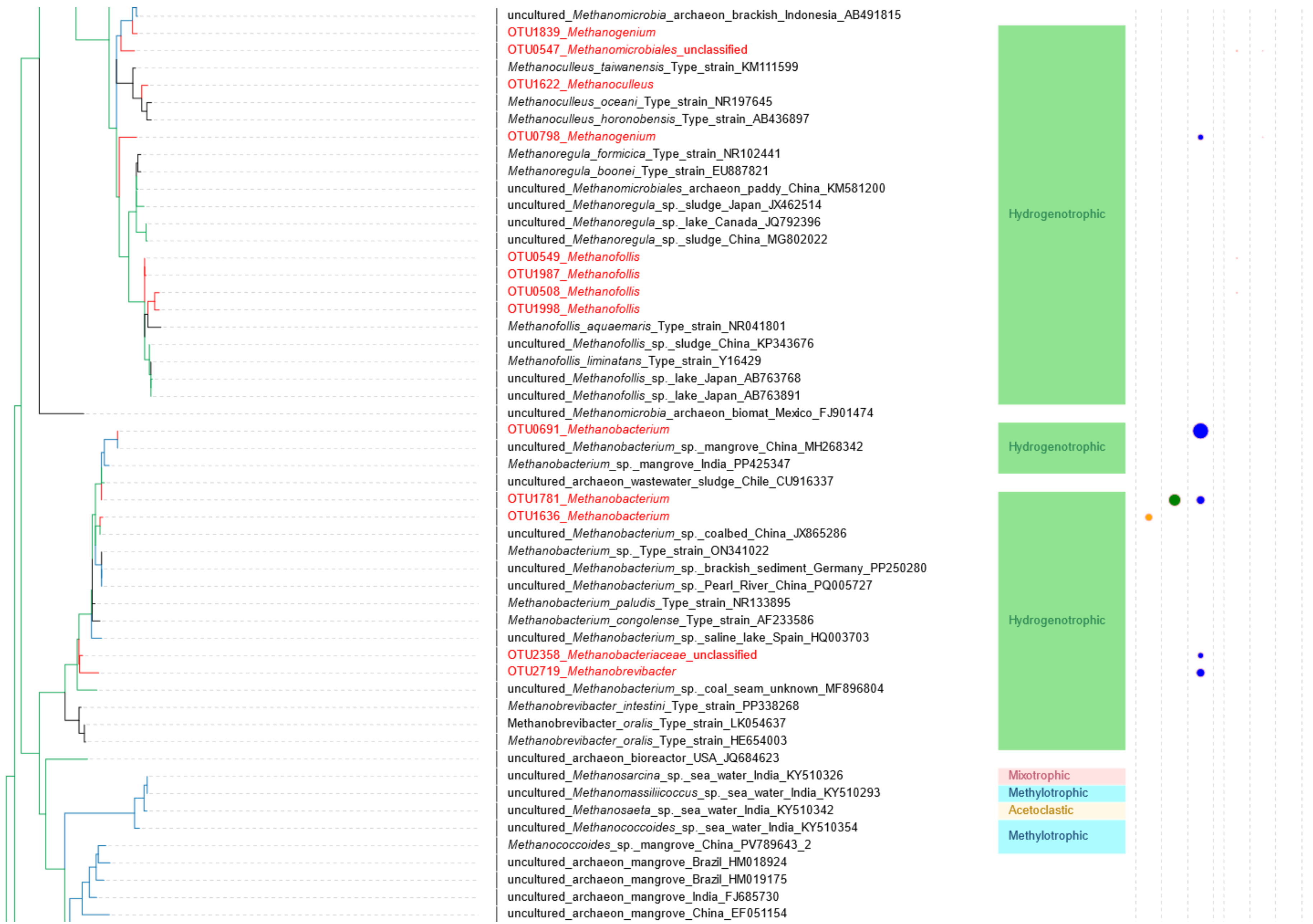


**Figure S6** continued

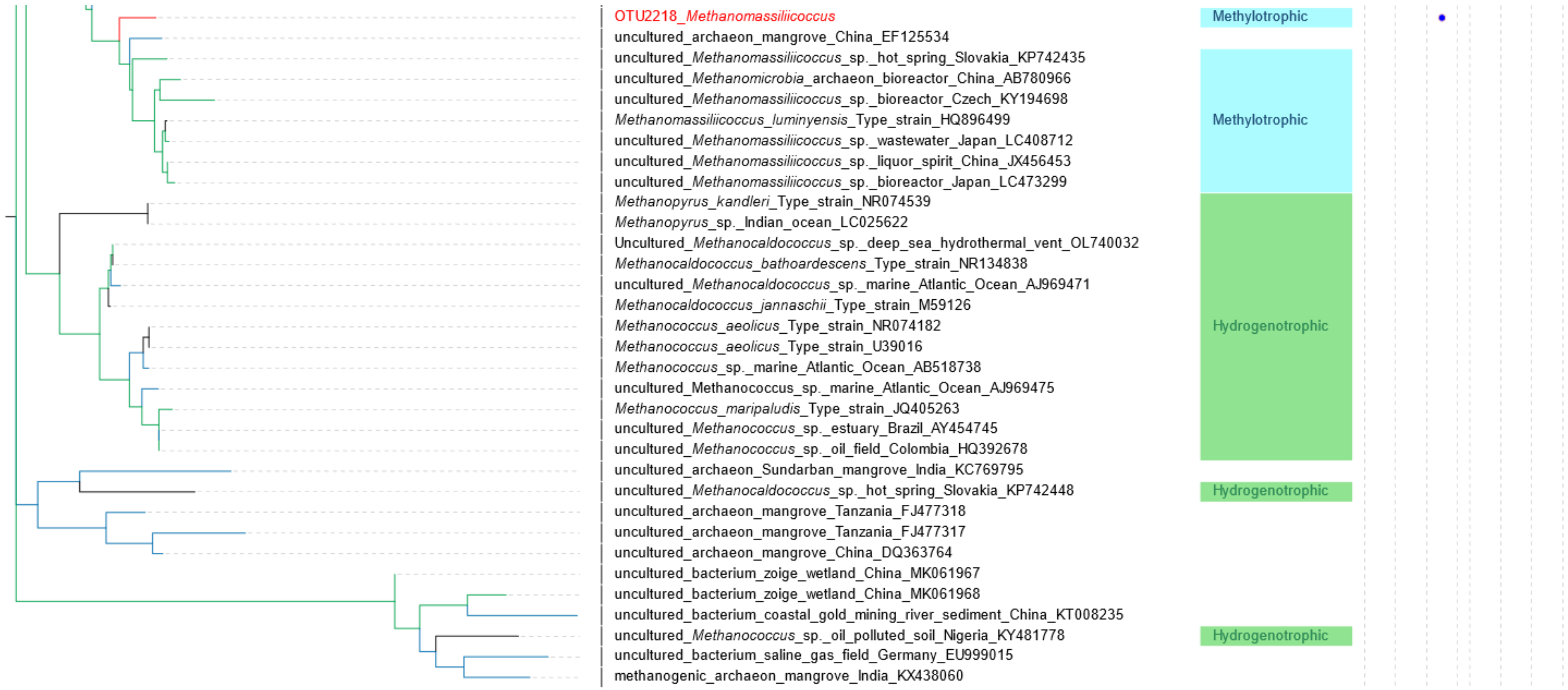


**Figure S6** continued

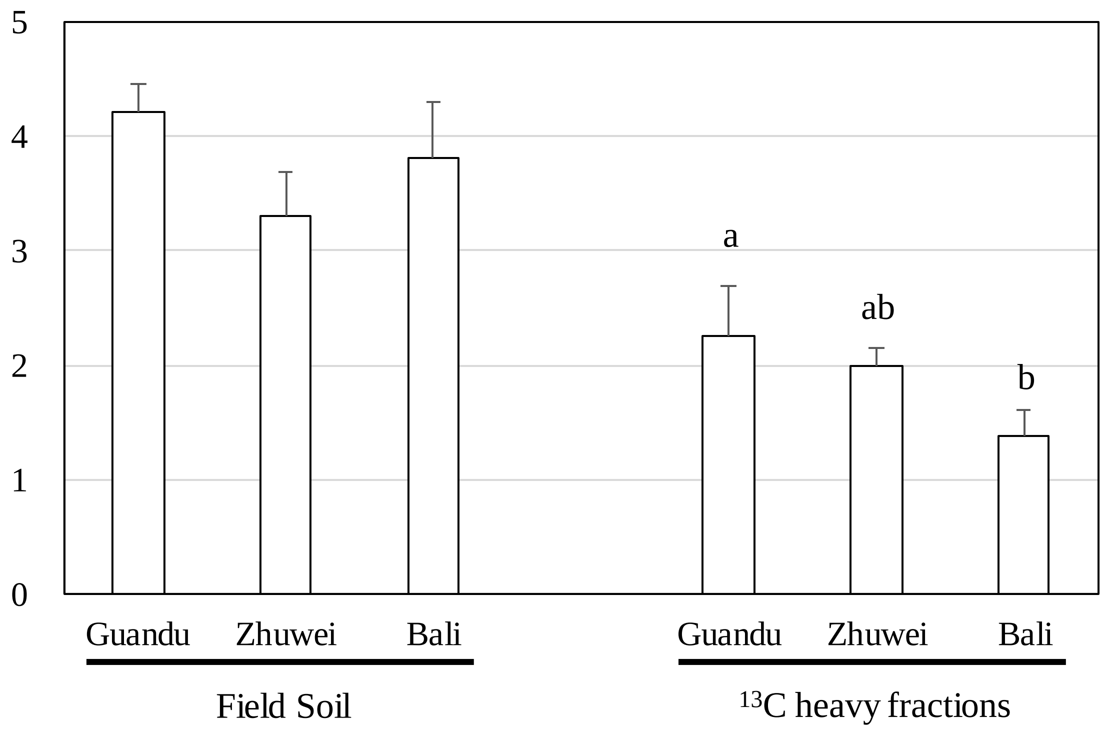


**Figure S7** Shannon diversity index of methanogenic communities based on *mcrA* gene sequences in field soils and $^{13}C$-methanol labeled heavy DNA fractions from three mangrove forests. Different letters indicate statistically significant differences among sites based on one-way ANOVA and Tukey's HSD test ($P < 0.05$).

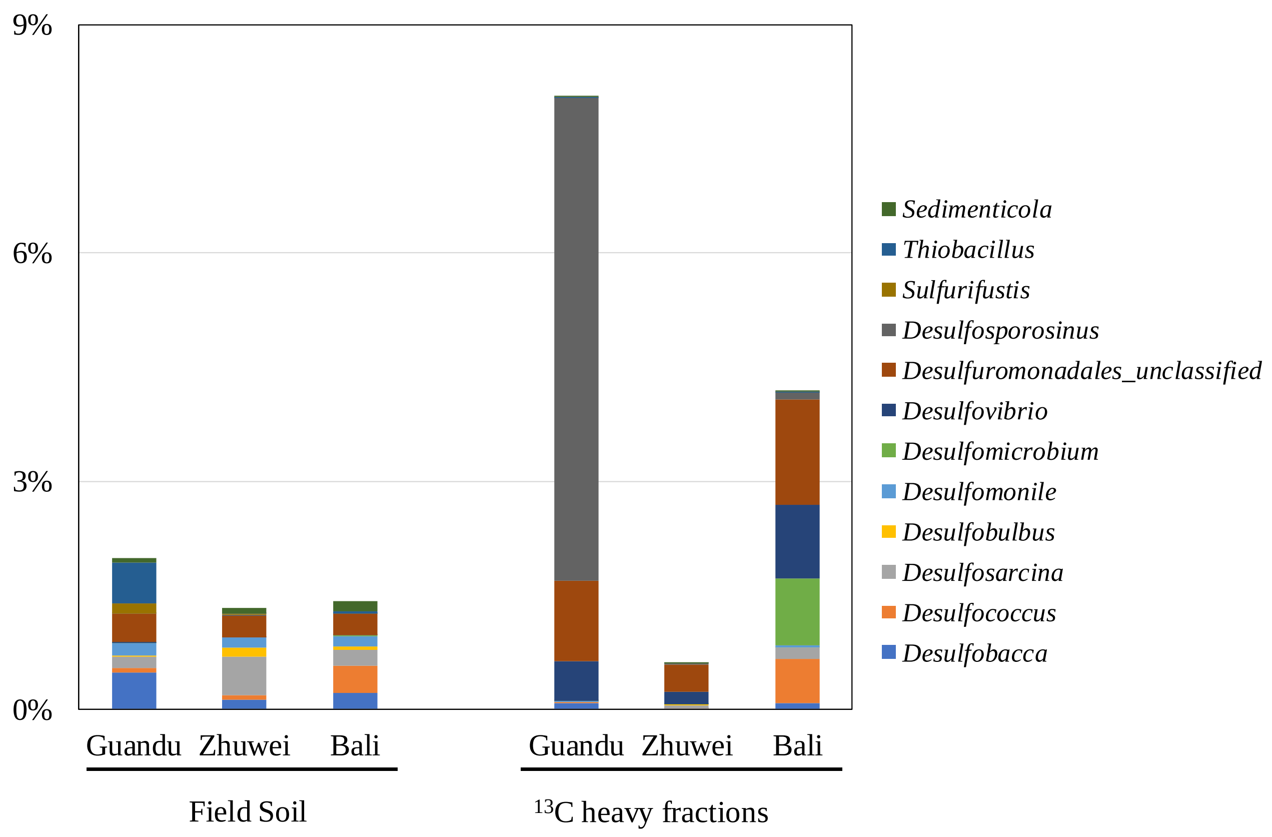


**Figure S8** Community composition of sulfate-reducing bacteria inferred from 16S rRNA gene amplicon data based on the KEGG PATHWAY database in field soils and $^{13}C$-methanol labeled heavy DNA fractions from three mangrove forests. The relative abundance shown on the y-axis represents the proportion of sequences assigned to sulfate-reducing bacteria relative to the total 16S rRNA gene sequences in each sample.

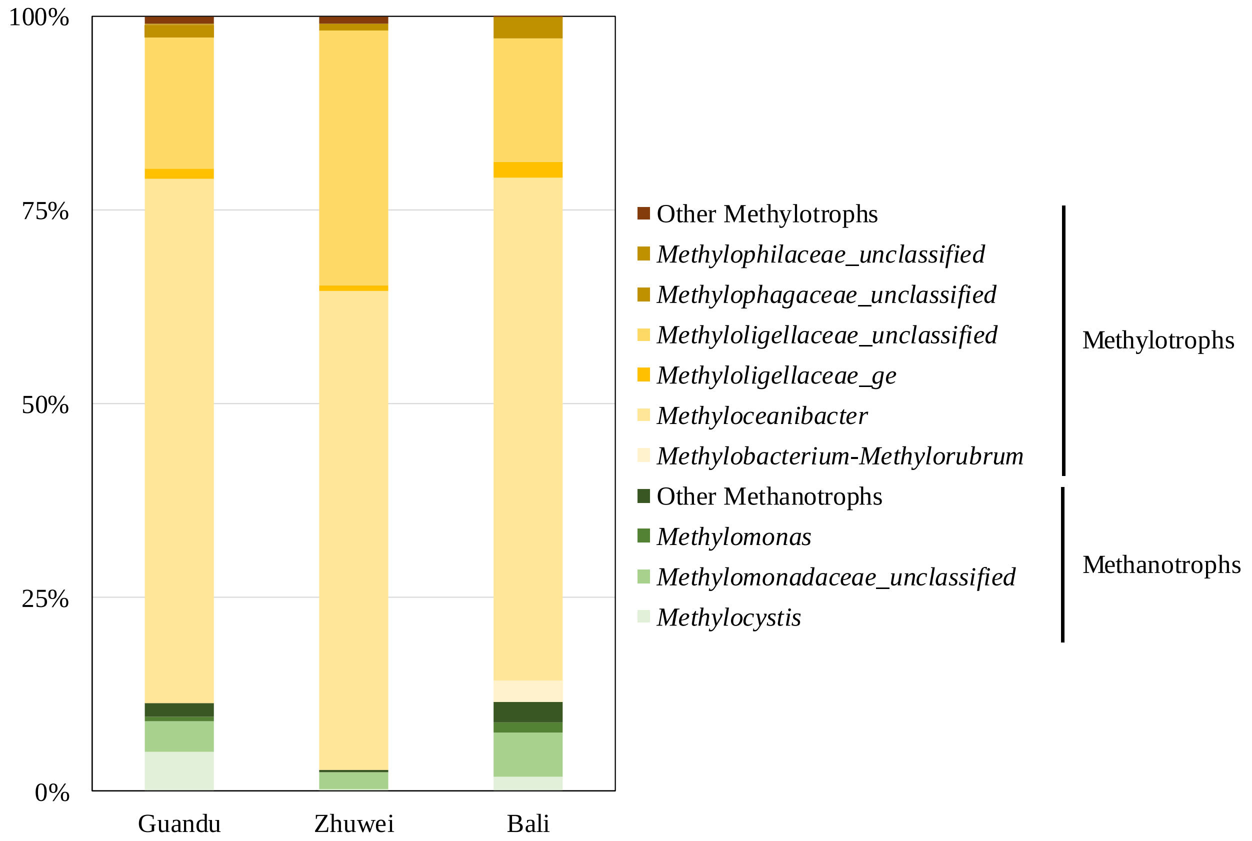


**Figure S9** Community composition of methylotrophic and methanotrophic communities inferred from 16S rRNA gene amplicon data based on the KEGG PATHWAY database in $^{13}$C-methanol labeled heavy DNA fractions from three mangrove forests. Relative abundance was calculated based on the total sequences classified as methylotrophic or aerobic methanotrophic lineages in each sample.
